# Effect of Delta and Omicron mutations on the RBD-SD1 domain of the Spike protein in SARS-CoV-2 and the Omicron mutations on RBD-ACE2 interface complex

**DOI:** 10.1101/2022.07.28.501901

**Authors:** Wai-Yim Ching, Puja Adhikari, Bahaa Jawad, Rudolf Podgornik

## Abstract

The receptor-binding domain (RBD) is the essential part in the Spike-protein (S-protein) of SARS-CoV-2 virus that directly binds to the human ACE2 receptor, making it a key target for many vaccines and therapies. Therefore, any mutations at this domain could affect the efficacy of these treatments as well as the viral-cell entry mechanism. We introduce *ab initio* DFT-based computational study that mainly focuses on two parts: (1) Mutations effects of both Delta and Omicron variants in the RBD-SD1 domain. (2) Impact of Omicron RBD mutations on the structure and properties of the RBD-ACE2 interface system. The in-depth analysis is based on the novel concept of *amino acid-amino acid bond pair units* (AABPU) that reveal the differences between the Delta and/or Omicron mutations and its corresponding wild-type strain in terms of the role played by non-local amino acid interactions, their 3D shapes and sizes, as well as contribution to hydrogen bonding and partial charge distributions. Our results also show that the interaction of Omicron RBD with ACE2 significantly increased its bonding between amino acids at the interface providing information on the implications of penetration of S-protein into ACE2, and thus offering a possible explanation for its high infectivity. Our findings enable us to present in more conspicuous atomic level detail the effect of specific mutations that may help in predicting and/or mitigating the next variant of concern.

## 1. Introduction

Back in late 2019, a novel coronavirus, the severe acute respiratory syndrome coronavirus 2 (SARS-CoV-2), was first identified as the causative agent for the virus disease and named COVID-19 by the World Health Organization (WHO)[1]. SARS-CoV-2 is continuously evolving due to genetic mutations or viral recombination during genome replication, resulting in emerging many Variants of Concern (VOCs) [2]. These VOCs significantly alter virus properties such as infectivity, transmissibility, antigenicity, and pathogenicity [3]. Further, some VOCs have the ability to evade natural or vaccine-induced immunity, decrease susceptibility to therapeutic agents, cause more severe disease, and spread faster [4]. As a result of these consequences, the number of confirmed cases and deaths is approaching to reach 550 million confirmed cases and more than 6.34 million deaths by July 5, 2022 [1].

Besides that, this terrible pandemic has a devastating effect on social, emotional, and economic consequences with no end in sight [5]. In response, intensive efforts have resulted in unprecedented success in the rapid development of COVID-19 vaccines, treatments, and diagnostics. The vaccination has proven to be effective in controlling the disease outbreak, especially for older patients and other more vulnerable groups [6]. Despite this success, the COVID-19 pandemic is far from over, with continuous emergence of new variants that are more dangerous and unpredictable. We all have to be prepared to fight a long battle in addressing new virus and find ways and means to deal with it.

In recent two years, WHO classified four variants as VOC including Alpha (B1.1.7), Beta (B.1.351), Gamma (P.1), and Delta (B.1.617.2) variant [7]. By the end of 2021, WHO also designated Omicron (B.1.1.529) as the most recent VOC [7]. The Omicron variant, B.1.529, is the most mutated SARS-CoV-2 variant, with 37 mutations in the spike protein (S-protein), 15 of which are in the receptor-binding domain (RBD), the primary target for vaccine and therapy development [8-12]. Within a few days, artificial intelligence (AI) modeling predicted [13] that the Omicron variant was 2.8 times more infectious than the Delta variant and had a nearly 90% chance of evading current vaccines. Subsequent experiments have confirmed Omicron’s high infectivity [14, 15], more vaccine breakthroughs [16, 17], and increased antibody escape rate [18-20]. Since January this year, several new Omicron sublineages have been continuously emerging, including BA.1, BA.1.1, BA.2, BA.2.12.1, BA.3, BA.4, and BA.5 [21, 22]. They share many mutations but also have significant differences. For example, BA.2 shares 32 mutations with BA.1 but differs by 28. Particularly, BA.2 has 16 mutations in the RBD of the S-protein, 12 of which are identical to those observed in BA.1, with the remaining four, S371F, T376A, D405N, and R408S, being unique. These differences suggest that there may be an effect on how these Omicron sublineages bind to ACE2 or monoclonal antibodies. On the other hand, Delta variant has only two RBD mutations, L452R and T478K. Interestingly, T478K was found in all Omicron sublineages, while these two Delta mutations appeared only in BA.4 and BA.5. Therefore, there is an urgent need to investigate how these mutations in RBD affect the structure and interatomic interactions with ACE2.

SARS-CoV-2 virus infection is a significant evolutionary event [23]. However, it is constantly evolving and mutating, resulting in the emergence of numerous major VOCs as discussed above. Therefore, recognizing mutational effects in circulating VOCs such as Delta and Omicron on the structure and dynamic processes of the S-protein is critical to gaining a fundamental understanding of how it maintains a strong interaction with ACE2, as well as establishing principles needed to guide the effective development of drugs or vaccines. These mutations depend on the locations and the nature of the substitution in the S-protein, which may lead to distinct alterations in their biological roles to influence viral fitness and pathogenicity [24, 25].

Numerous remarkable achievements have been made in VOCs research such as determining their viral genomic sequences, resolving the three-dimensional (3D) structures of the S-protein and the interface RBD-ACE2 systems, as well as characterizing their viral entry mechanisms [26-33]. These achievements would not have been possible without many focused efforts utilizing various methodologies. For instance, sequence homology tools [34, 35] have been used on various aspects of mutations related to existing variants and their subvariants. There are also reports providing a comprehensive survey of experimental and clinical studies on SARS-CoV-2 pathologies and the human proteins associated with these pathologies [36-39]. Other studies show that binding free energy (BFE) between the RBD in S-protein and ACE2 is proportional to the viral infectivity [13, 40, 41]. Apparently, natural selection favoring the more infectious variants is part of the fundamental law of biology that governs SARS-CoV-2 transmission and evolution [42, 43], including the occurrence of Alpha, Beta, Gamma, Delta, and Omicron variants.

More recently, there was a flurry of research activity [44, 45] based on artificial intelligence (AI) and machine learning (ML) techniques, such as AlphaFold [46] and RoseTTAFold [47]. These algorithms can predict the 3D shape of proteins by exploiting the relationship between experimentally determined residue sequences in the deposited databases of various sources, and thereby lead to the next stage of understanding and predicting the protein folding. These are indeed wonderful and great achievements in biomedical research. Nevertheless, it has been pointed out that the protein-folding problem is not yet completely solved [48, 49]. Structural prediction, no matter how accurate they are, lack specific details on interatomic interactions which is the domain of quantum mechanics [50, 51]. Prominent British Science writer Philip Ball [52] pointed out that in fact subjects such as replication, mutation, and selection are still not much explored at the level of single molecules or amino acids (AAs) [53].

As of today, there are many computational approaches and strategies to address the above shortcomings. They all have strong supporters and specific merits, but they also have noticeable drawbacks and limitations. Classical molecular dynamics (MD) based on putative carefully designed force fields has received a lot of traction [54-56], while other attempts were based upon quantum chemical calculations or their modifications, such as on the density functional theory (DFT) [57, 58] with specific modifications to accomplish different goals. The most common shortcoming of the latter approaches is still the limited molecular size that can be accommodated and the demand for prohibitive computational resources, making these calculations impractical. Nonetheless, much progress has been made in recent years by compromises, balancing the time needed and the cost involved. One example is *ab initio* fragment molecular orbital (FMO) approach, which divides a large biomolecule into small fragments and performs molecular orbital (MO) calculations on each fragment and its dimers to determine the properties of the entire system [59]. FMO has been used to investigate SARS-CoV-2 S-protein interactions with ACE2 or antibodies [60-62]. We are also at the frontier of this grand challenge by using a *divide-and-conquer strategy (DCS)* that allows us to apply the DFT calculations to large systems, such as the S-protein, by concentrating on each of its individual structural domains. With this approach, we have been able to run a single DFT computation for each specific domain with up to 5000 atoms [63-69]. In-depth analysis of interatomic interactions centered on the novel concept of *amino acid-amino acid bond pair units* (AABPU) disclosed important insights into the mutational effects of the S-proteins. This is also the main goal of this paper.

In this feature article, we present a comprehensive study of mutation consequences on the structure and properties of the RBD-SD1 domain in the S-protein of SARS-CoV-2 and the RBD-ACE2 interface. Part-1 of the paper focuses on the mutation effects of Delta and Omicron variants in the RBD-SD1 segment which contain 2 and 16 mutations respectively. Part-2 shifts to the impact of the Omicron mutations on the interface of the RBD-ACE2 complex. These results are based on large quantum mechanical calculations at atomic scale (discussed in **Section S1** of supplementary materials (SM)), using the novel concept of AABPU as critical structural units. The combination of these results provides a promising pathway by using purely computational means to mitigate any future variants of concern and in fighting the pandemonic of the century.

## 2. Model specification

### 2.1 RBD-SD1 in S-protein

The present work consists of two parts. The first part is described here, and it focuses on the receptor-binding domain (RBD) with subdomain 1 (SD1) of the SARS-CoV-2 S-protein, which is referred to as the RBD-SD1 model and shown in **Figure 1**. The initial structure for the region of RBD-SD1 was obtained from Woo et al (PDB ID: 6VSB) [70], which originated from Wrapp et al [41].

**Figure 1.**
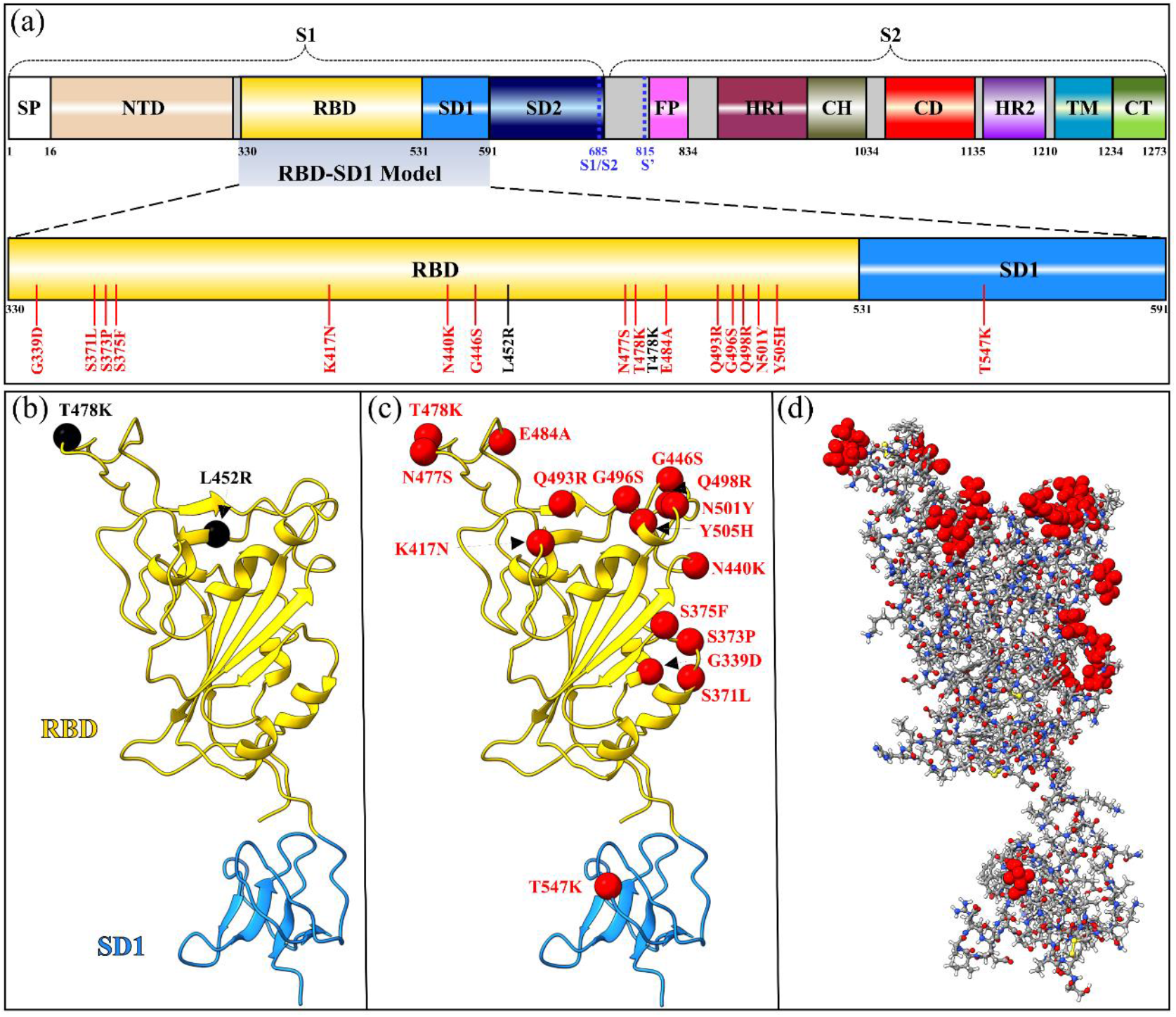
RBD-SD1 models in both Delta and Omicron variants. (a) Schematic depiction of S-protein primary structure divided into domains with cleavage sites S1/S2 and S2’. The RBD-SD1 regions with all their DV and OV mutations are highlighted and zoomed in. (b) The ribbon structure of RBD-SD1 model of DV with two mutations marked by the black circles. (c) The RBD-SD1 model of BA.1 OV with 16 mutations marked by the red circles. (d) The ball and stick model in (c) with 16 OV mutations marked by the red circles.

In this part we compared Delta variant (DV) and Omicron variant (OV) with the Wild type (WT). There are total of 4059, 4072, and 4123 atoms in WT, DV, and OV model, respectively.

From the Woo et al structure, we chose chain A of S-protein in its up confirmation. Sequence 330-591 was selected from the S-protein for RBD-SD1 (6VSB_1_2_1) model [70]. The glycans associated with the PDB are removed. Furthermore, hydrogen atoms were added using the Leap module with ff14SB force field in the AMBER package [71]. For RBD-SD1 DV, the two L452R and T478K mutations, shown in **Figure 1 (a)** and (**b**), were generated using Dunbrack backbone-dependent rotamer library as implemented in the UCSF Chimera package [72].

RBD-SD1 OV model, has sixteen mutations: G339D, S371L, S373P, S375F, K417N, N440K, G446S, S477N, T478K, E484A, Q493R, G496S, Q498R, N501Y, Y505H, and T547K as shown in **Figure 1 (a)** and (**c**). The conformations of K417N and N501Y were modeled using PDB ID 7V80 [73] while T478K using 7ORA [74]. The remaining thirteen OV mutations were modeled using the conformations with the highest probabilities from the Dunbrack backbone-dependent rotamer library [75].

### 2.2 RBD-ACE2 Interface Complex

The second part of the present work focuses on the interactions at the interface between RBD and a portion of ACE2 for WT and OV. The structures of the interfaces were obtained from the PDB ID 6M0J [76] for the WT and from PDB ID 7WBP[77] for the OV. The model constructed for the calculation of RBD-ACE2 is displayed in **Figure 2**. Amino acids were included from the sequence S19-I88 and G319-T365 in the ACE2 region [66, 68] and from sequence T333-G526 in RBD. The entire model has 311 amino acids (194 in RBD and 117 in ACE2). There are 4817 and 4873 atoms in the WT and OV interface models, respectively. In this OV RBD-ACE2 model there are only fifteen mutations since SD1 is not included. All the fifteen OV mutations are marked with red color in **Figure 2**. Hydrogen atoms were added using the Leap module with ff14SB force field in the AMBER package [71].

**Figure 2.**
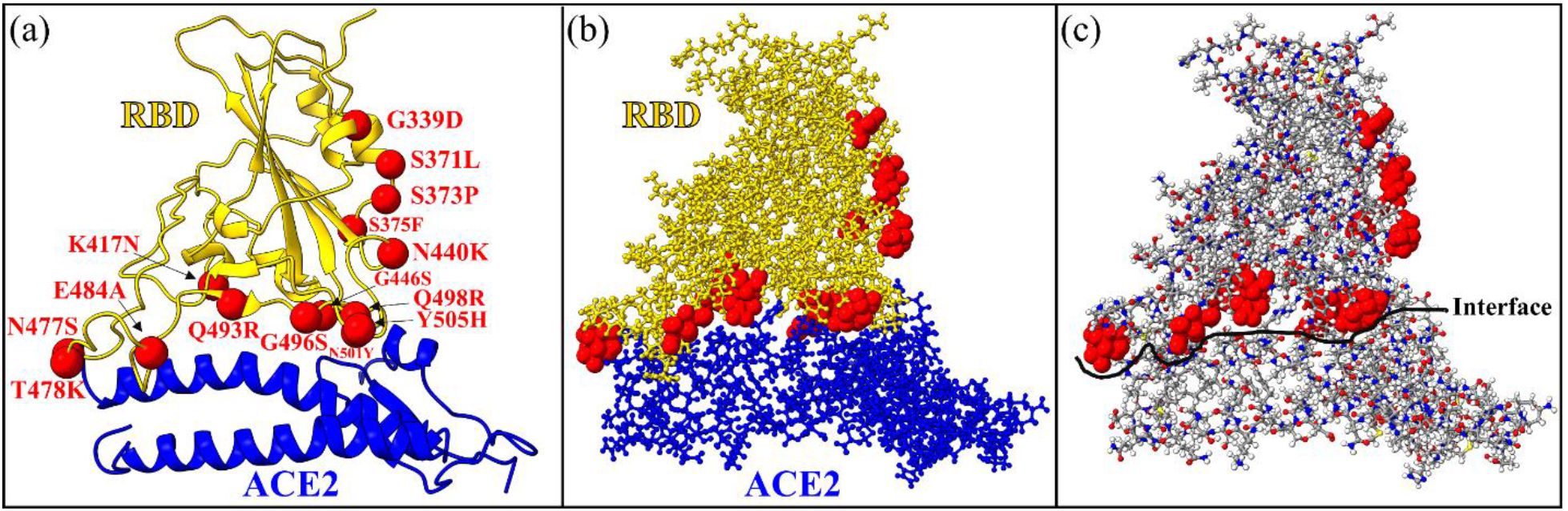
The RBD-ACE2 interface model. (a) Ribbon structure showing the interface between RBD and the segment of ACE2. Mutated AAs of the RBD in OV are marked by red sphere. (b) The ball and stick structure of the same model as in (a). (c) alternative version of (b) with more details of the atoms. Grey: C, red: O, blue: N, and white: H. In the WT interface model, there are 2993 atoms in RBD and 1824 atoms in ACE2 segment with a total of 4817 atoms. In the OV interface model, there are 3049 atoms in RBD and 1824 atoms in ACE2 segment for a total of 4873 atoms.

## 3. Results: Part 1

### 3.1 Changes in the RBD-SD1 S-protein due to Delta and Omicron mutations

In this study, two well-known packages based on density functional theory (DFT) have been used: the Vienna ab initio Simulations package (VASP) [78] and the orthogonalized linear combination of atomic orbital OLCAO) technique [79]. These DFT calculations can provide a lot of key parameters useful in probing mutational impacts as detailed in SM.

In **Table 1**, we list the result of AAPBU analysis for the 18 mutations sites from WT among which 2 sites are for DV and 16 sites are for OV. WT T478 and OV K478 are listed twice since they are in both DV and OV. For the details of the definitions of the AABP, nearest neighbor NN-AAPB, non-local neighbor NL-AAPB, contribution of hydrogen bonds to AABP (HB), number of non-local AAs, as well as partial charge (PC), consult the SM. Partial charge for the AABPU (PC*) will be presented in more detail in **Section 3.4**. Here it should be mentioned that the values present in **Table 1** are estimated for the entire AABPU, not for a single amino acid site (see SM).

**Table 1.**
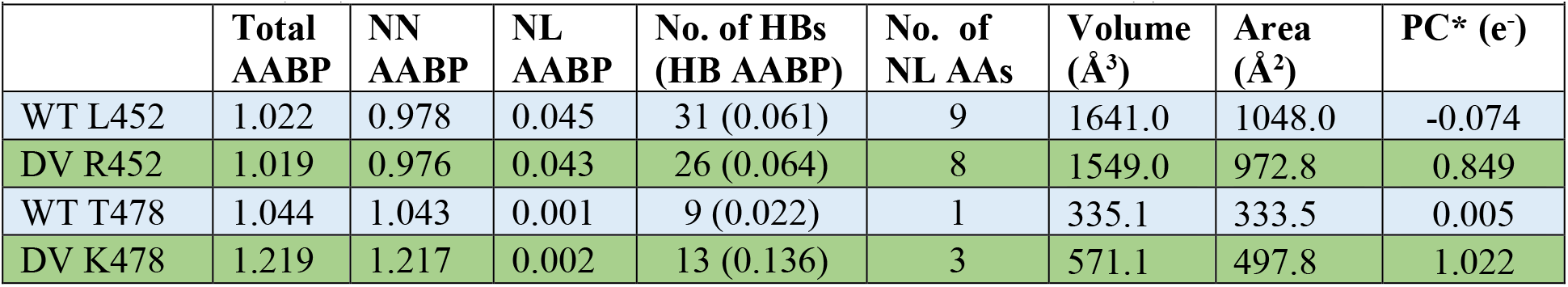
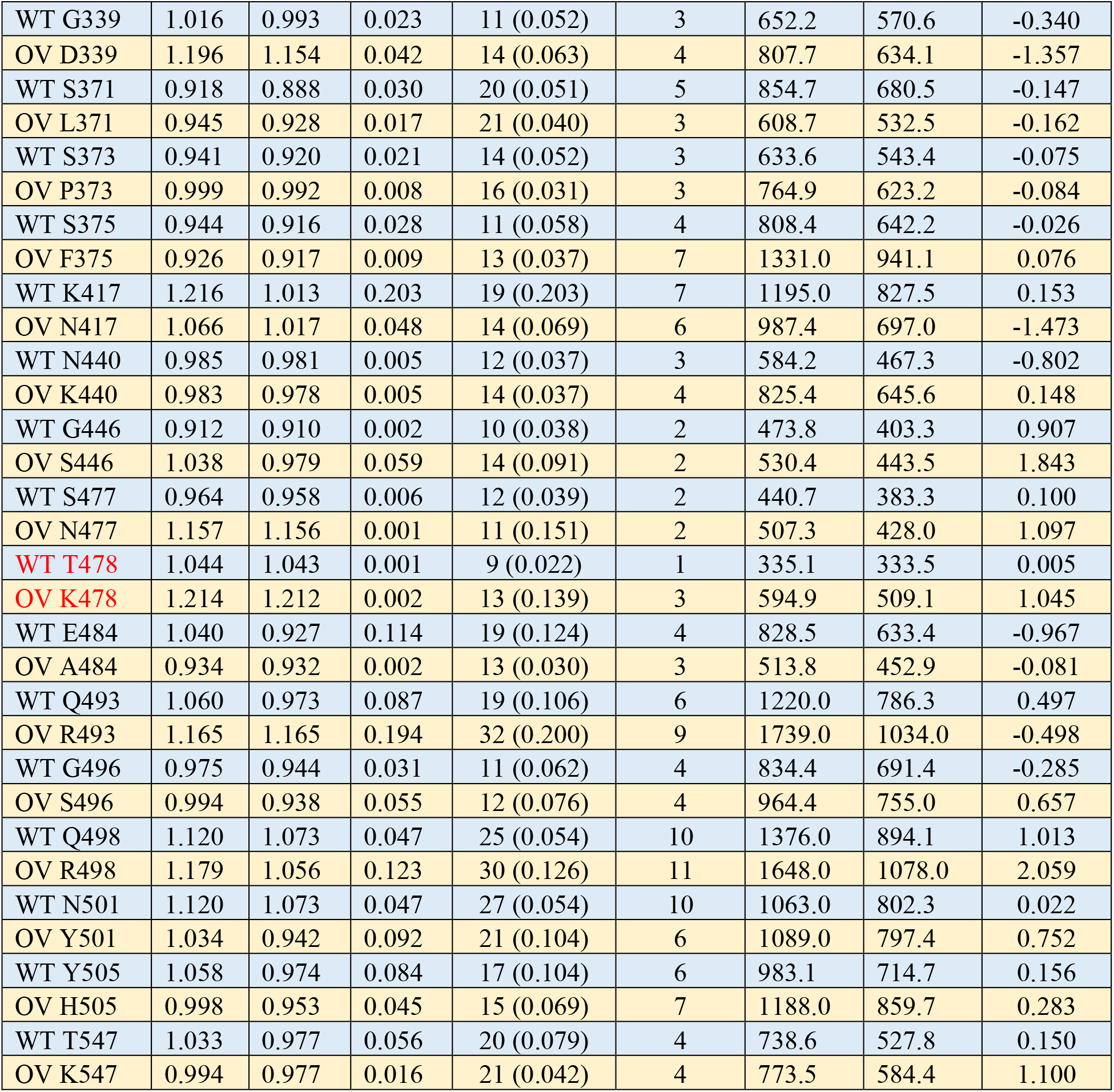
Comparison of AABP units between wild type (WT), top 2 for Delta variant (DV) and 16 Omicron variant (OV) in RBD-SD1 domain. AABP is in unit of electrons (e).

From **Table 1** and based on plots in **Figure 3** and **Figure S1**, we have extracted the following detailed description of the mutations which are as a rule missing in the characterization available in the literature. A succinct list of these observations is listed below. We focus on the relative quantitative differences as presented in **Table 1**, underlying the spectacular variety of mutation changes.

**Figure 3.**
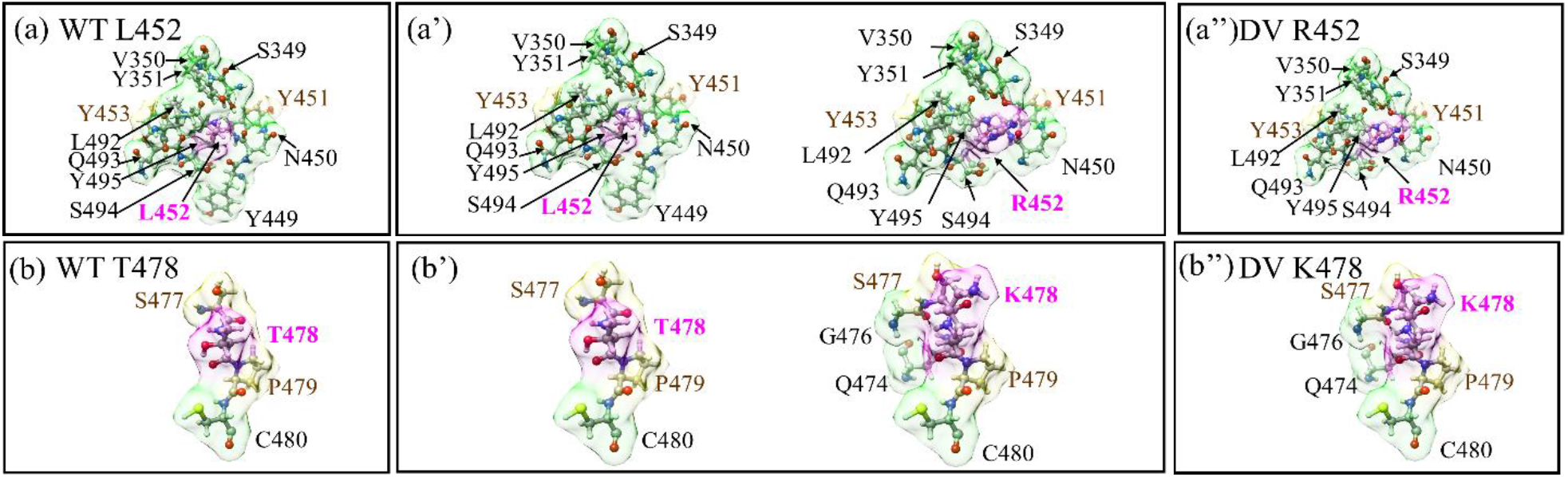
Comparison of changes in the shape of AABPU for the 2 DV mutations with respect to their WT sites. and (a’) are shown in the fixed scale for WT and DV for site 452 for comparison. (a) and (a’’) are shown in the real scale. Similarly (b), (b’), and (b’’) shows for the site 478.

1. Thel argest AABP in WT is K417 (1.216 e^-^) and mutated one is DV K478 (1.219 e^-^).
2. The smallest AABP in WT is G446 (0.912 e^-^) and mutated one is OV A484 (0.934 e^-^).
3. The largest NN-AABP in WT is in both N501 and Q498 (1.073 e^-^) and mutated one is DV K478 (1.217 e^-^).
4. The smallest NN-AABP in WT is S371 (0.888 e^-^) and mutated one is OV F375 (0.917 e^-^).
5. The largest NL-AABP in WT is K417 (0.203 e^-^) and mutated one is OV R493 (0.194 e^-^).
6. The smallest NL-AABP in WT is T478 (0.001e^-^) and mutated one is OV N477 (0.001 e^-^).
7. The largest contribution from hydrogen bonds (HB) to total AABP in WT is K417 (0.203 e^-^) and in mutated one is OV R493 (0.200 e^-^). This finding is similar to the one observed for the largest NL-AABP, indicating that HB plays a dominant role in NL-AABP.
8. The smallest contribution from HB to total AABP in WT is T478 (0.022 e^-^) and in mutated one is OV A484 (0.030 e^-^).
9. The overall comparison in number of HBs in the 18 mutation sites are further shown in **Figure S2** for simplicity. OV R493 has highest difference of HBs after mutation. The total difference in number of HB in 16 OV mutations sites (OV-WT) is 18 and in 2 DV mutations sites (DV-WT) is -1, inducting substantial change in the intramolecular HB distributions of OV RBD-SD1.
10. The largest number of NL AAs in WT is 10 at Q498 and N501 and for mutated one is 11 at R498 (OV).
11. The smallest number of NL AAs in WT is 1 at T478 and mutated one is 2 at S446 (OV) and N477 (OV).
12. The largest volume of AABPU is L452 (1641.0 Å^3^) in WT and OV R493 in mutated one (1739.0 Å^3^).
13. The smallest volume of AABPU is T478 (335.1 Å^3^) in WT and OV N477 in mutated one (507.3 Å^3^).
14. The largest and smallest surface area of AABPU are correlated with their volume as expected.
15. We notice there are some mutated AAs closer together (clustering effect) forming large and small groups. The 4 clusters are (S371L, S373P, S375F), (N440K, G446S), (S477N, T478K, E484A), (Q493R, G496S, Q498R, N501Y, Y505H).

In **Figure 3**, we compare the mutational changes in the shape of AABPU for the first 2 mutations in DV (WT L452, WT T478 to R452 and K478). They depend on the scale of the plot used. Fixed scale makes their boxes the same. Real scale reveals the change in the shape of AABPU. It can be seen that mutation L452R reduce its volume whereas mutation T478K increase its volume as listed in **Table 1**. Similar figures for 16 OV mutations are shown in **Figure S1**.

In **Figure 4**, we display each component of the AABPU data in **Table 1** in bar graph form except the PC which will be discussed in **Section 3.4. Figure 4 (a)** displays the total AABP values for the 18 mutations and **Figure 4 (b)** to **(f)** displays NN AABP, NL AABP, AABP from HB, Volume, and surface area of each AABP unit, respectively. It can be summarized briefly as follows:

**Figure 4.**
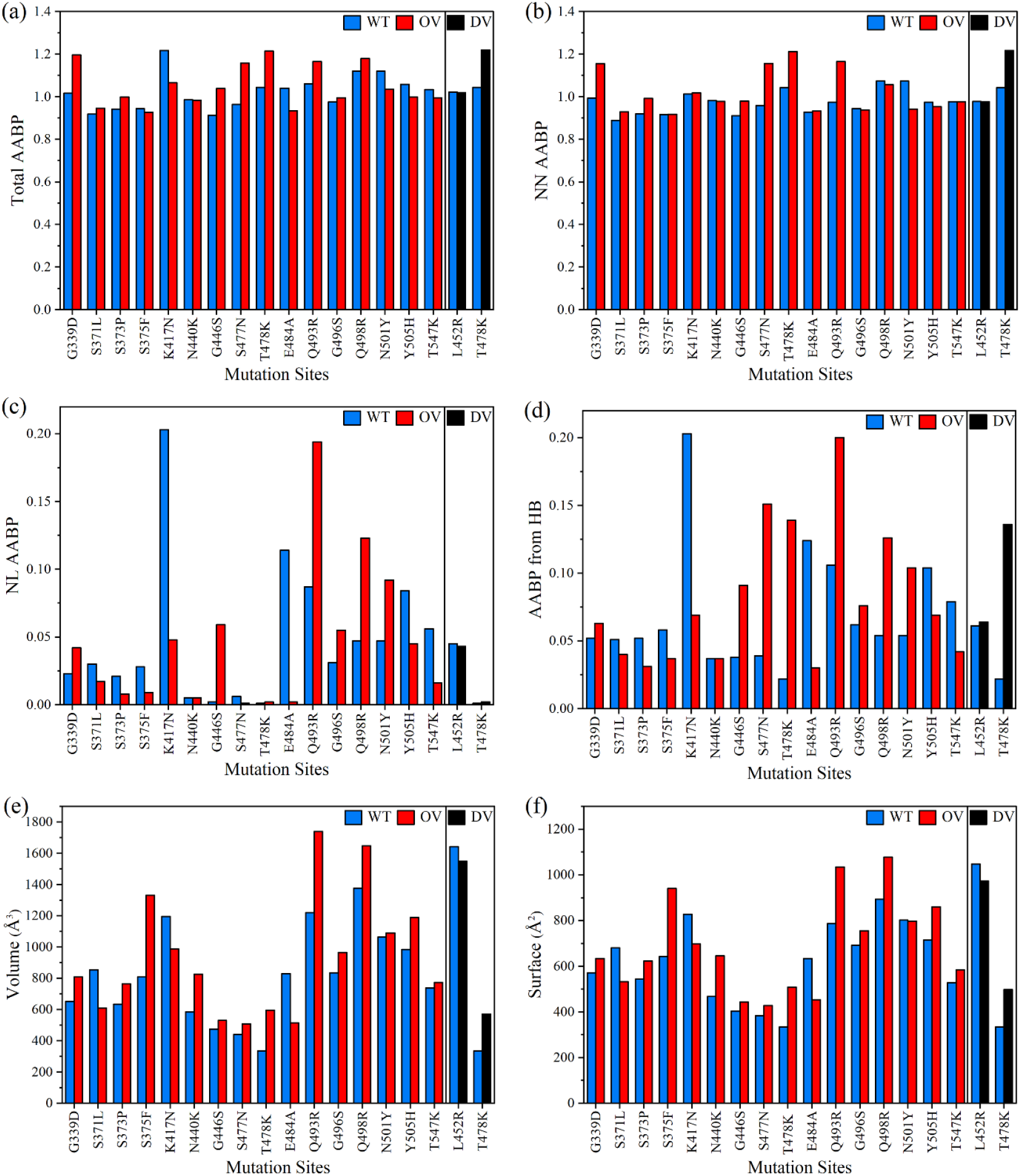
Comparison of 2 DV and 16 OV mutations with their corresponding WT sites of RBD-SD1 model in term of. (a) Total AABP, (b) NN AABP, (c) NL AABP, (d) AABP from HB, (e) Volume, (f) Surface area.

1. Total AABP and NN AABP differs only slightly accentuating the importance of using sequence of AAs in proteins and their fundamental analysis.
2. The contributions from NL AABP and hydrogen bonding (HB) to AABP are non-negligible.
3. Depending on the type of AAs substitution, location of the mutation site, and interatomic interactions, the AABP changes can either increase or decrease.
4. Large changes in the volumes of the AABP units upon mutation. Except for DV L452R and OV S371L, K417N, and E484A, all other mutations result in increase in the volume of these units.
5. Changes in surface areas of AAPBU correlates with the change in volume. Slight difference in trends reflects the change in the shape of some of AABPU.

### 3.2 Electronic Structure of Delta and Omicron RBD-SD1

In condensed matter physics, the electronic structure of a material, usually a crystalline material, is generally presented and interpreted in terms of the total density of states (TDOS)—a plot showing the distribution of the calculated energy eigenvalues as the number of states per energy level. The occupied portion or the valence band (VB) and unoccupied portion or the conduction band (CB) are separated by a and gap E_g_ for insulators or a Fermi level (E_F_) with no gaps for metals. In quantum chemistry, the top of VB and bottom of CB are respectively called HOMO (highest occupied molecular orbitals) or LUMO (lowest occupied molecular orbitals) separated by a HOMO-LUMO gap. In complex biomolecules similar data can be presented since *ab initio* calculation using OLCAO method give all the energy eigenvalues. The TDOS can be resolved into partial DOS (PDOS) for each atom or a group of atoms in a specific structural unit which are usually well-defined even for very complex crystals. In complex biomolecules such as in the present work, the decomposition of TDOS into PDOS is far more challenging but can still be done if the partial structural units are clearly specified. This provides insightful information for a deeper level of understanding in their electronic structure.

Figure 5. shows the plot of TDOS for the RBD-SD1 unit for the three models WT, DV and OV with slightly different total number of atoms of 4059, 4072 and 4123, respectively. They have only minor differences in TDOS since for such large systems, the changes in atomic configurations are small. Their HOMO-LUMO gaps are about 1.68 eV. The TDOS is obtained from a huge number of energy eigenvalues for all atoms interacting in each model. The TDOS in **Figure 5** and be resolved into 18 PDOS. They are discussed in SM in **Section S2** (Partial density of states (PDOS) for WT, DV and OV in RBD-SD1) and **Figure S3**.

**Figure 5.**
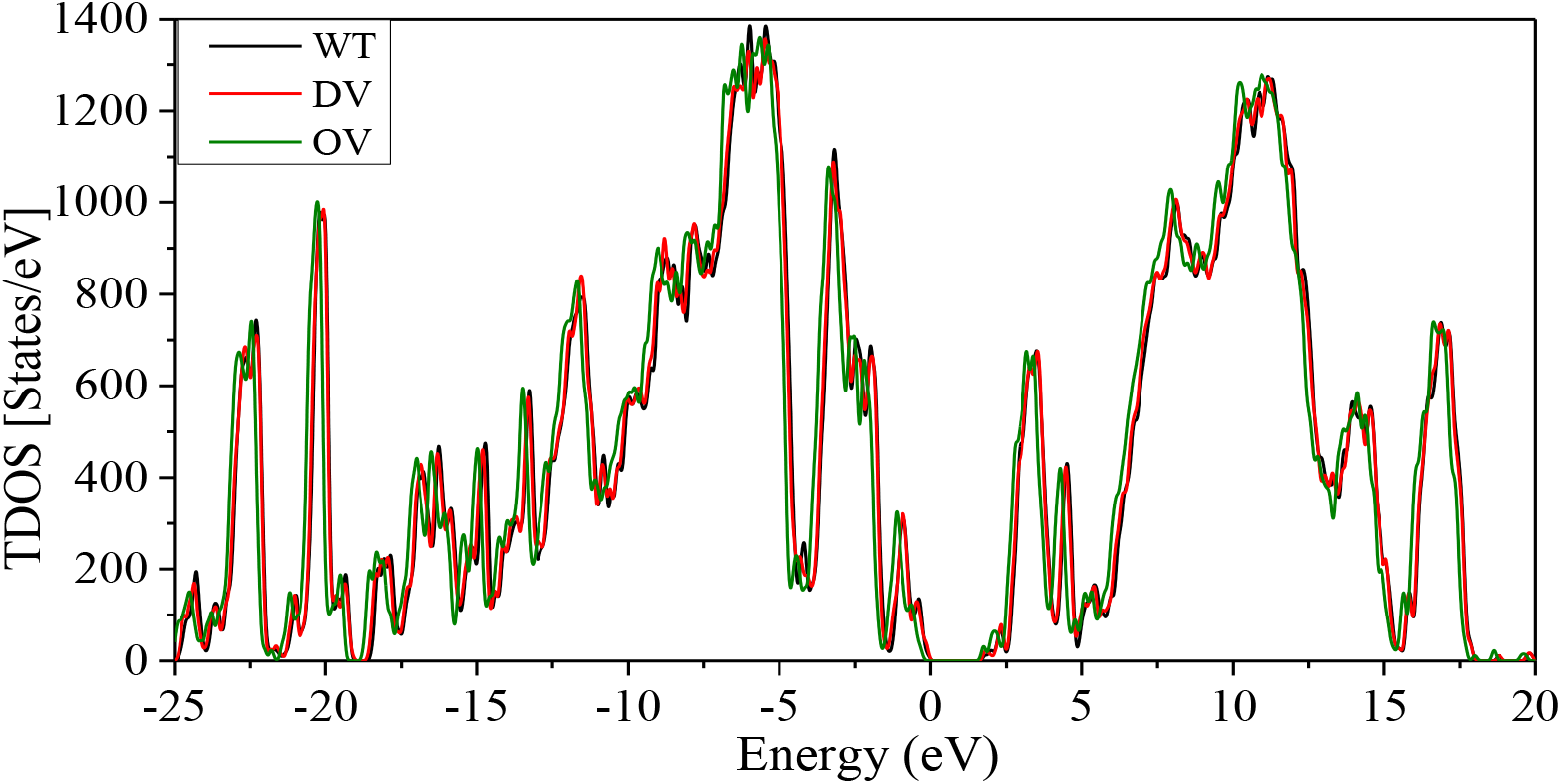
Total density of states (TDOS) for the RDB-SD1 model in WT, DV and OV. HOMO-LUMO gap is 1.68 eV.

### 3.3 Interatomic Bonding in Delta and Omicron RBD-SD1

The data for bond order (BO) vs. bond length (BL) distribution for all the atomic pairs in RBD-SD1 domain in WT, DV, and OV models are displayed in **Figure 6**. There are 20299 data points for WT, 20410 data points for DV, and 20709 data points for OV for a total of 61,418 data points. The figure shows the atomic-scale interactions for all atoms in RBD-SD1 models within the BL range of up to 4.5 Å as well as the effect of mutations for each pair, demonstrating the detailed atomic-scale level that quantum chemical calculation can achieve. Of particular importance is the distribution of Hydrogen bonding (HB) which is ubiquitous. HB is probably the most important bonding in biomolecules but is seldom discussed in detail. The quantification of HB network has been previous described solely based on the HB in water, that forms a HB network [80]. It is usually assumed that HBs are weak, but they are of pivotal importance in any biological system especially in proteins. **Figure 6 (b)** shows that HB in RBD-SD1 can occur first at the BL close to 1.51 Å and with BO value close to 0.139 e^-^. They can be affected by mutations in both DV and OV but predominately in the OV. The change in number of HBs in 18 sites of WT in comparison to DV, and OV are shown in **Figure S2**. As already listed in the HB contribution to the total AABP value in **Section 3.1** and **Table 1**. The HB data in RBD-SD1 for WT, DV and OV are briefly summarized in **Table S1**. There are total of 12,142 HBs (4021 for WT, 4037 for DV, and 4084 for OV) out of 61,418 data points. Again, the strongest HB shown in **Figure 6 (b)** first appears at BL of 1.51 Å with BO of 0.139 e^-^ and then continues to other BL/BO combinations in **Figure 6 (c)** and **(d)** with gradually decreased BO values. One can also discern on the unusual HBs of different types mostly with tetrahedrally bonded C [81] and we plan to make a separate and detailed analysis of these HBs in RBD-SD1 and will be presented in a separate publication [82].

**Figure 6.**
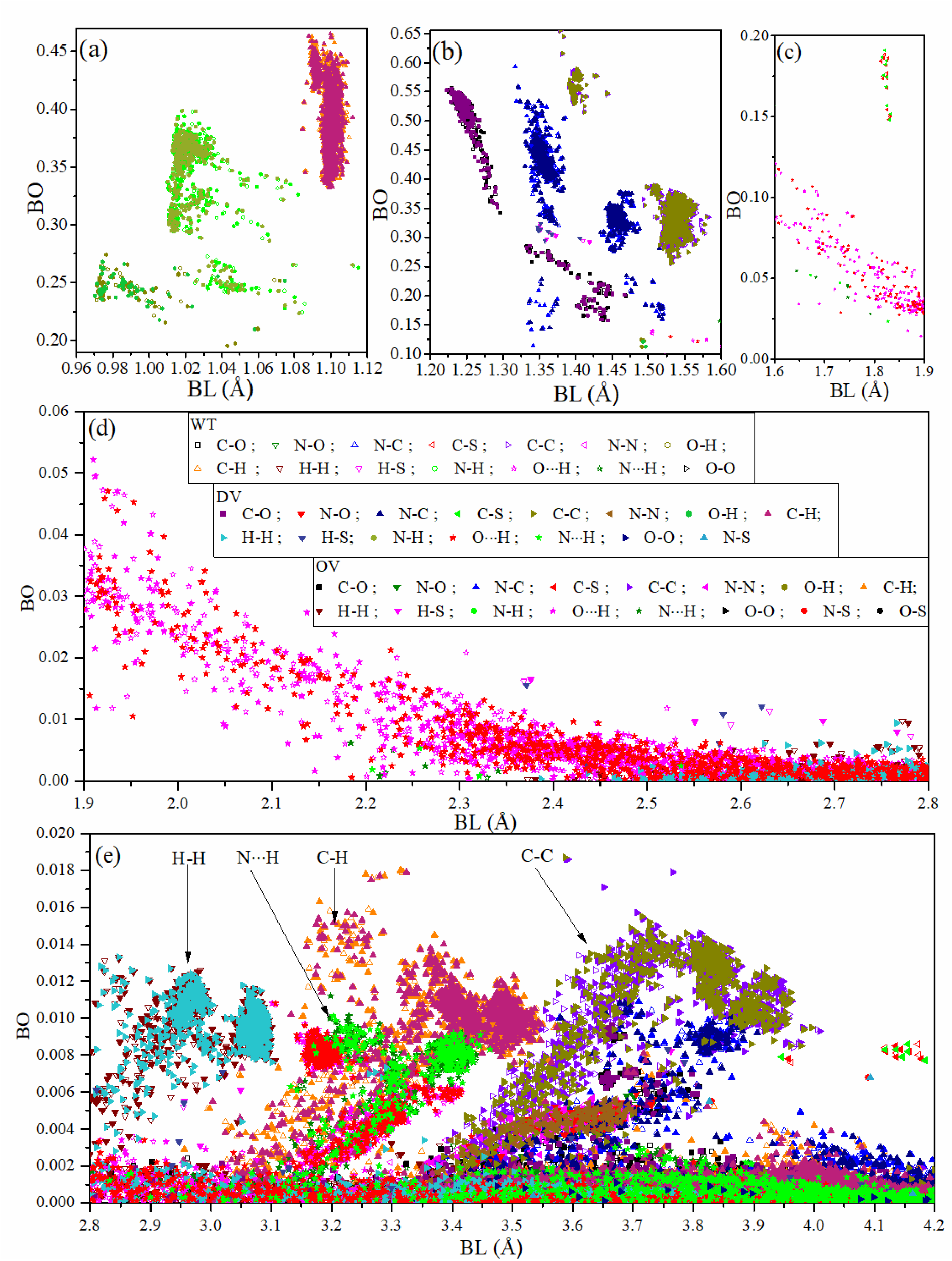
BO vs BL distribution for the RBD-SD1 in WT, DV, and OV: (a) 0.96 - 1.12 Å, (b) 1.20 – 1.60 Å, (c) 1.60– 1.90 Å, (d) 1.90 – 2.80 Å, and (e) 2.8-4.2 Å. Each of the 61,418 data point is designated in the inset in (b). Specific bonding groups at large BL are indicated by vertical arrows in (c).

The complex mixture of different types of bonds in **Figure 6 (a)** to **(e)** is identified by their ranges of BL for easy recognition. The symbols for these data points are depicted in the inset in **Figure 6 (b)** in three groups: WT, DV and OV. They are succinctly summarized as follows:

1. **Figure 6 (a)**: From the BL range 0.96 Å to 1.08 Å, the data points are labeled as WT O-H, DV O-H, OV O-H, WT N-H, DV N-H, OV N-H. These are the first group of covalent bonds. From BL range 1.08 Å to 1.12 Å, the data points are WT C-H, DV C-H, OV C-H. These are the second group of stronger covalent bonds with higher BO.
2. **Figure 6 (b)**: Overlapping groups between different type of atoms in AAs. The group with BL of 1.22 Å to 1.46 Å consists of the C-O covalent bonds in three cases (WT, DV, and OV). For the group from 1.33 Å to 1.52 Å, they consist of covalent bonds between N and C, N and O, H and S and some O-H bonds in WT, DV, and OV. The other two overlapping groups from 1.39 Å to 1.45 Å and 1.50 Å to 1.58 Å labeled as WT C-C, DV C-C, OV C-C are basically covalent bond between C atoms from same or different AAs but at longer distances of separation. Finally, HBs start to appear between 1.51 Å to 1.60 Å.
3. **Figure 6 (c)**: There are two distinct groups. From BL 1.60 Å to 1.90 Å, are weaker HBs. From the narrow range of 1.83 Å to 1.84 Å, the data are from C-S bond with relatively larger BO values than HBs (WT C-S, OV C-S, DV C-S).
4. **Figure 6 (d)**: Weaker HBs (N…H, O…H) in the range from 1.90 Å to 2.80 Å in WT, DV, OV. From 2.37 Å to 2.80 Å, second nearest neighbor (NN) bonds with S and also remote H-H interactions start to appear (WT H-S, DV H-S, OV H-S, WT H-H, DV H-H, OV H-H). These and other very weak remote second NN H-S bonds extending beyond 2.8 Å will be depicted in **Figure 6 (e)**.
5. **Figure 6 (e)**: More HBs present in this region. Clustering of specific 2^nd^ NN bond groups are marked separately. These are all very weak interactions with low BO values, but they are ubiquitous and collectively make non negligible contribution in proteins.

The above issues on specific types of interatomic bonding are seldom discussed in the literature, especially the very weak bonds. Only the full atomic scale *ab initio* calculation interpreted in the framework of the AABPs, as detailed in this work, gives access to these data.

### 3.4 Partial Charge Distribution of Delta and Omicron RBD-SD1

Partial charge (PC) is a key parameter in electrostatic potential of a biomolecule crucial for predicting the overall long-range intermolecular interactions. Our *ab initio* calculated PCs are potentially useful for computing this electrostatic interaction using Delphi software, for example [66, 83]. It could also be used to improve the accuracy of the PCs used in the most of MD simulations. In this study, we have two types of PC. One is PC for each specific AAs (PC^AA^), and another is PC for the entire AABPU unit (PC*). PC* is obtained adding up PC^AA^ of all AAs involved in the AABPU. **Figure 7 (a)** shows PC* distribution of the AABPU listed in **Table 1** for the 18 mutations (2 DV and 16 OV). 10 out of 16 in OV (62.5%) have positive PC* whereas the two DV mutations are both positive. The two DV mutations L452R and T478K have huge increase from near zero PC* in WT to close to 1 e. Among the 16 OV mutations, only 3 of them are shifted toward negative PC* (G339D, K417N, Q493R) with the last 2 flip form positive to negative PC*. Additionally, the three mutations located outside of RBM (S371L, S373P, and S375F) together with Y505H exhibit no substantial change in their PC*. Finally, G446S, S477N, T478K, Q498R, N501Y, and T547K OV mutations cause a change in the charge distribution toward a more positively charged state, as with N440K and G496S mutations, but this time switching from negative to positive PC*. Similarly, E484A is also alter the PC* toward positive state but remains in a negative state.

**Figure 7.**
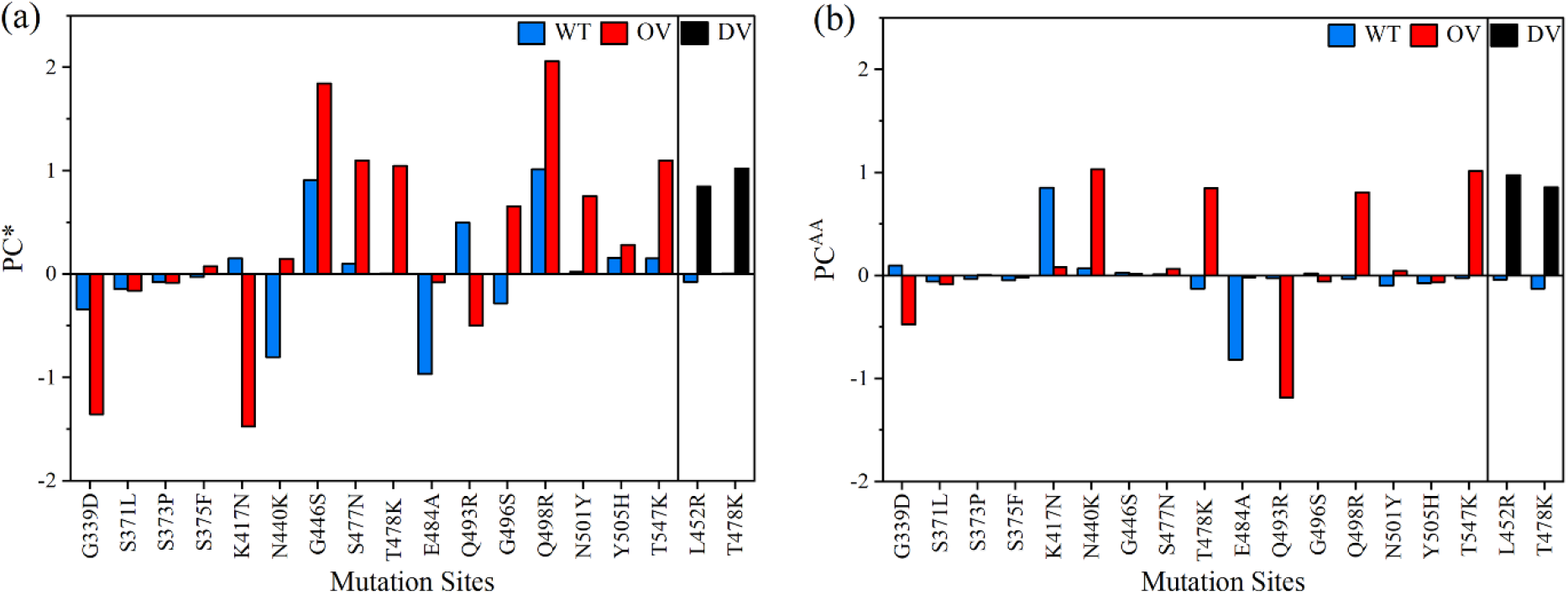
(a) Partial charge per AABPU (PC*) and (b) partial charge distribution of each AAs (PC^AA^) in unit of electron (e) at 18 mutation sites of RBD-SD1.

The partial charge is emerging as an important proxy for the quantification of mutational drift of the different VOC in particular as it seems to increase in a steady progression towards the Omicron variant [68, 84]. The appropriate quantification of the partial charge can only be obtained from *ab initio* DFT calculation and has profound implications since all major SARS-CoV-2 variants have been accumulating positive charges in solvent-exposed regions of the S-protein, especially its ACE2-binding sites or along the RBD epitopes that are targeted by many therapeutic antibodies. More specific, the accumulation of positive mutations in Omicron RBD results an increase in the total charge density of the S-protein, facilitating the recognition process with the negative charge of ACE2 (as will discuss later in Part 2) or participating in immune evasion [8, 32, 85].

In **Figure 7(b)**, we show the PC^AA^ of the key AA at 18 mutation sites and compare with PC* for the whole unit of AABPU in **Table 1**. They basically mimic the results in **Figure 7(a)**, especially for the 2 DVs. The only exceptions are the two OV in G339D, K417N with negative PC*. Some of the differences between **Figure 7(a)** and **7(b)** could be attributed to the fact that we performed our simulation at neutral pH, while the pH impact could play an important role in the partial charge distribution and regulation. More detail investigation in this respect is clearly beyond the scope of current investigation.

## 4. Results: Part 2

### 4.1 Differences and similarities between unbound Omicron RBD and bound RBD with ACE2

Again, the RBD-ACE2 interface model, as shown in **Figure 2**, contains 311 AAs (194 in RBD and 117 in ACE2), with the Omicron RBD having 15 mutations, one less compared to the unbound RBD-SD1 model in Part 1 since the subdomain SD1 is not included (see **Section 2.2**).

Data from **Table 2** together with **Figure 8** and **Figure S4** show considerable changes when compared with **Table 1 (Figure 4** and **Figure S2)** in Part 1. Most of these changes are due to the mutations in AABPU that result in stronger interactions between RBD with ACE2. Here are some observations comparing the RBD-ACE2 and RBD-SD1 models: (a) There are different AABP values (in **Table 1** and **Table 2**) because of the differences in number of NL interacting AAs. However, 10 out of 15 AAs exhibit similar trend of total AABP. i.e., either total AABP of WT is higher, or total AABP of OV is higher in both tables. This difference in remaining 5 AAs could be traced to the fact that there is an extra SD1 in Part 1 and RBD’s interaction with ACE2 in Part 2. (b) K417 (WT) has highest NL AABP in both RBD-ACE2 and RBD-SD1 models. Similarly, Q493, E484, and Y505 from WT and R493, and R498 from OV have higher NL AABP in both cases. (c) R493 from OV has significant contributions in both the models. (d) The difference (OV-WT) in total number of HBs for 15 mutations sites is 17 for RBD-SD1 whereas 8 for RBD-ACE2 due to dissimilarity in number of NL AAs. (e) There are differences in volume and surface values for AABPU, since the number of NL AAs are quite different. (f) Most of the volume and surface area of AABPU increase after mutation in both models. However, there is decrease in volume in four cases (K417N, G446S, E484A and Y505H) of RBD-ACE2 and three cases (S371L, K417N, and E484A) of RBD-SD1 model. The change in the surface area follows the change in volume closely but not exactly since the shape of each AABPU alters due to difference in NN and NL AAs.

**Table 2.** Comparison of AABP units for the 15 OV mutations in RBD-ACE2 complex with their corresponding WT sites. AABP is in unit of electrons (e).

|  |  |  |  |  |  |  |  |  |
| --- | --- | --- | --- | --- | --- | --- | --- | --- |
|  | **Total AABP** | **NN AABP** | **NL AABP** | **No. of HBs (HB AABP)** | **No. of NL AAs** | **Volume (Å<sup>3</sup>)** | **Area (Å<sup>2</sup>)** | **PC\* (e)** |
| WT G339 | 1.032 | 0.989 | 0.043 | 14 (0.070) | 5 | 1628.0 | 1288.0 | -0.532 |
| OV D339 | 1.111 | 1.002 | 0.109 | 18 (0.126) | 5 | 1629.0 | 1334.0 | -1.313 |
| WT S371 | 1.027 | 0.969 | 0.059 | 17 (0.077) | 5 | 857.3 | 680.1 | -0.121 |
| OV L371 | 0.927 | 0.904 | 0.023 | 16 (0.045) | 6 | 1274.0 | 989.3 | 0.034 |
| WT S373 | 1.009 | 0.932 | 0.077 | 11 (0.100) | 3 | 650.1 | 530.7 | 0.024 |
| OV P373 | 1.010 | 1.008 | 0.003 | 14 (0.021) | 3 | 791.2 | 619.7 | 0.053 |
| WT S375 | 1.028 | 0.998 | 0.030 | 11 (0.110) | 6 | 1116.0 | 858.0 | 0.826 |
| OV F375 | 1.135 | 1.081 | 0.055 | 11 (0.200) | 6 | 1177.0 | 896.6 | 1.005 |
| WT K417 | 1.388 | 1.014 | 0.374 | 24 (0.123) | 9 | 1434.0 | 1035.0 | -0.744 |
| OV N417 | 1.106 | 1.028 | 0.077 | 16 (0.095) | 8 | 1347.0 | 908.9 | -0.875 |
| WT N440 | 0.923 | 0.919 | 0.004 | 11 (0.035) | 2 | 496.2 | 414.3 | -0.845 |
| OV K440 | 1.198 | 0.919 | 0.280 | 12 (0.040) | 3 | 636.3 | 587.0 | -0.894 |
| WT G446 | 1.024 | 0.972 | 0.052 | 17 (0.075) | 4 | 770.5 | 616.8 | 0.724 |
| OV S446 | 0.995 | 0.937 | 0.058 | 15 (0.083) | 3 | 695.3 | 549.7 | 1.297 |
| WT S477 | 0.965 | 0.952 | 0.013 | 12 (0.044) | 2 | 444.6 | 387.4 | 0.126 |
| OV N477 | 1.147 | 0.938 | 0.209 | 17 (0.216) | 5 | 821.7 | 704.2 | 1.694 |
| WT T478 | 1.050 | 1.048 | 0.002 | 13 (0.023) | 3 | 616.9 | 527.6 | -0.087 |
| OV K478 | 1.078 | 1.014 | 0.064 | 18 (0.022) | 4 | 741.2 | 667.7 | 1.189 |
| WT E484 | 1.170 | 0.928 | 0.242 | 22 (0.248) | 4 | 808.9 | 672.9 | -0.132 |
| OV A484 | 0.931 | 0.926 | 0.005 | 13 (0.033) | 2 | 447.3 | 393.9 | -0.096 |
| WT Q493 | 1.213 | 0.966 | 0.248 | 28 (0.256) | 8 | 1459.0 | 955.1 | -0.315 |
| OV R493 | 1.348 | 1.071 | 0.277 | 32 (0.324) | 11 | 2021.0 | 1217.0 | 1.314 |
| WT G496 | 1.043 | 0.976 | 0.067 | 15 (0.089) | 6 | 1210.0 | 1033.0 | -0.218 |
| OV S496 | 1.012 | 0.928 | 0.084 | 22 (0.103) | 7 | 1245.0 | 844.7 | -0.266 |
| WT Q498 | 1.277 | 1.083 | 0.194 | 35 (0.185) | 14 | 2071.0 | 1344.0 | 1.654 |
| OV R498 | 1.291 | 1.052 | 0.239 | 39 (0.222) | 14 | 2059.0 | 1232.0 | 0.943 |
| WT N501 | 1.134 | 0.948 | 0.186 | 29 (0.183) | 9 | 1699.0 | 1261.0 | -0.255 |
| OV Y501 | 1.029 | 0.946 | 0.083 | 20 (0.088) | 9 | 1912.0 | 1319.0 | 0.539 |
| WT Y505 | 1.341 | 1.002 | 0.339 | 23 (0.128) | 10 | 1773.0 | 1265.0 | 0.018 |
| OV H505 | 1.106 | 0.975 | 0.131 | 27 (0.142) | 9 | 1628.0 | 1161.0 | 0.287 |

**Figure 8.**
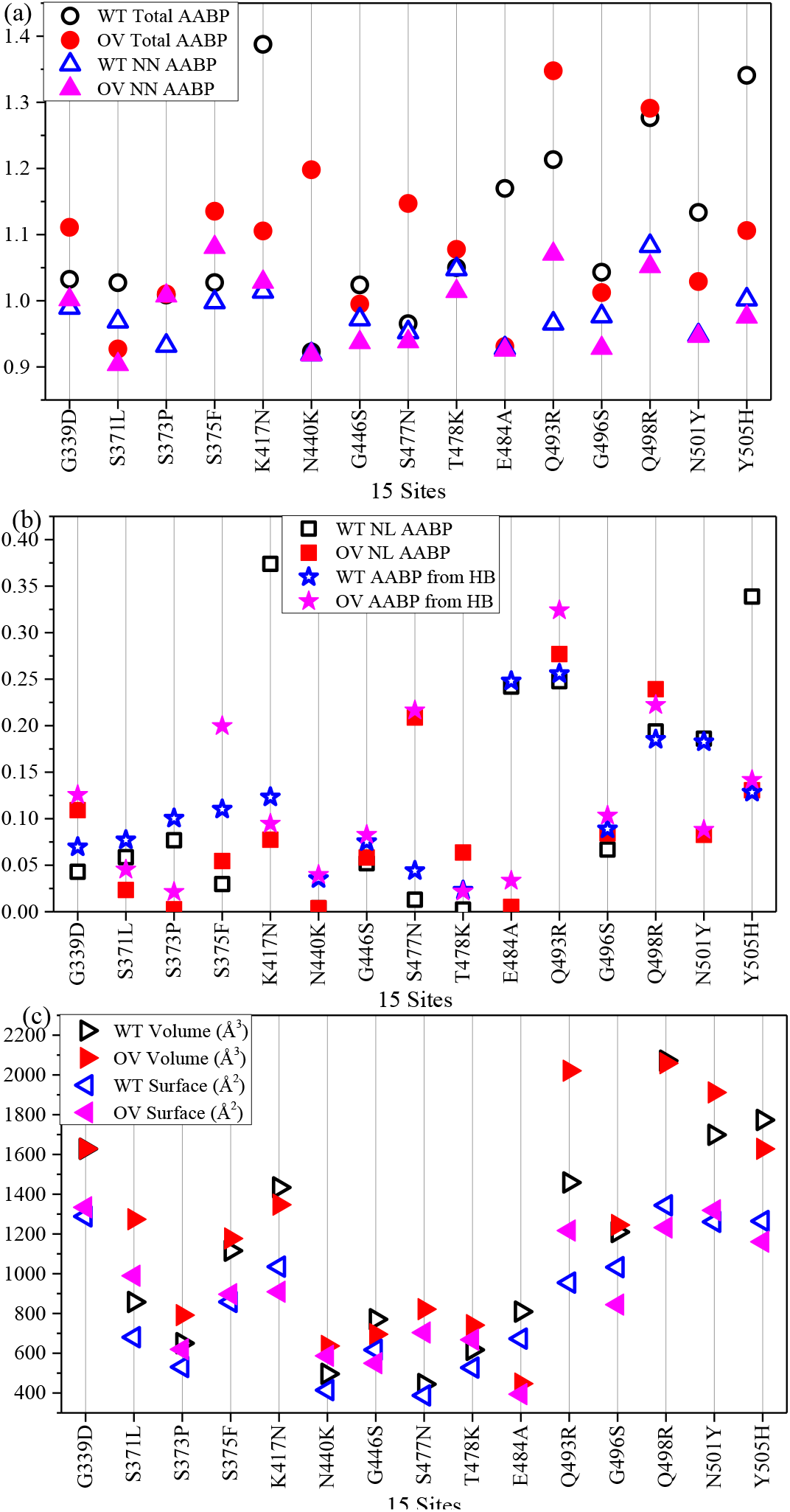
Comparison of 15 OV mutations with their corresponding WT sites of RBD-ACE2 complex in term of (a) Total AABP and NN AABP, (b) NL AABP and AABP from HB, and (c) volume and surface for WT and OV.

Figure 8. shows the complexity of the changes in the AABPU of the RBD at the interface with ACE2. Similar figures such as **Figure S1** in Part 1 can be plotted but are not included here. Additional figures involving the ACE2 part of the interface will be presented later in **Section 4.2** and **4.3**.

### 4.2 Properties and Interactions at RBD-ACE2 Interface

The total density of states (TDOS) for the interface model RBD-ACE2 has been calculated in the usual manner as in part 1. The TDOS is resolved into PDOS for RBD and ACE2 in **Figure 9** with each panel contains the WT and OV parts. As expected, these PDOS plots are very close to each other with only minor differences in peak structures in all biomolecules. The HOMO-LUMO gaps for RBD and ACE2 are 1.83 eV and 1.47 eV in RBD and ACE2, respectively. Based on HOMO-LUMO gap the Fermi level can be analyzed and modified to prepare materials such as sensors [86].

**Figure 9.**
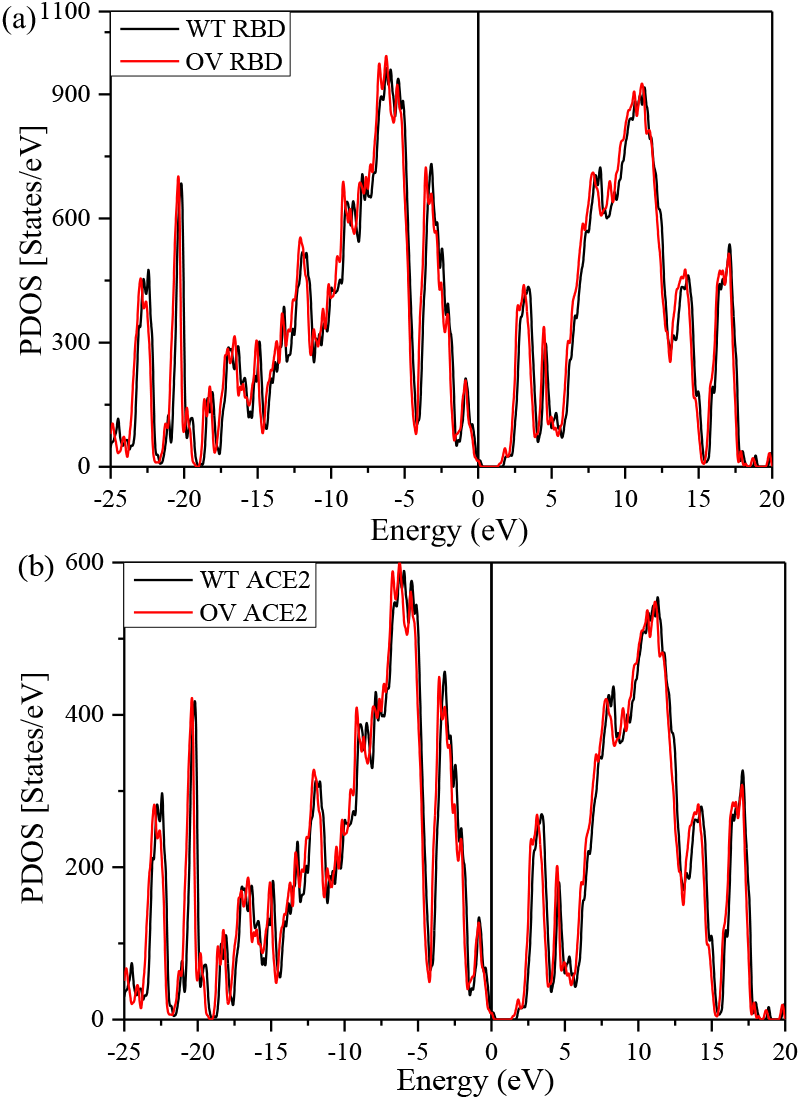
PDOS for (a) RBD and (b) ACE2. Each panel contains both WT and OV models.

Interaction between RBD and ACE2 is illustrated in **Figure 10** with the AABP values. **Figure 10 (a)** and **(b)** focus on the bonding between mutated and unmutated AAs of RBD and ACE2, comparing their AABP values, respectively. Adding the AABP values gives us the total AABP of 1.33e and 1.46e for WT and OV respectively. Hence, the OV has stronger binding with ACE2 than WT. The present interface calculation is superior to our past calculation on RBM-ACE2 [68] since the current calculation includes entire RBD.

**Figure 10.**
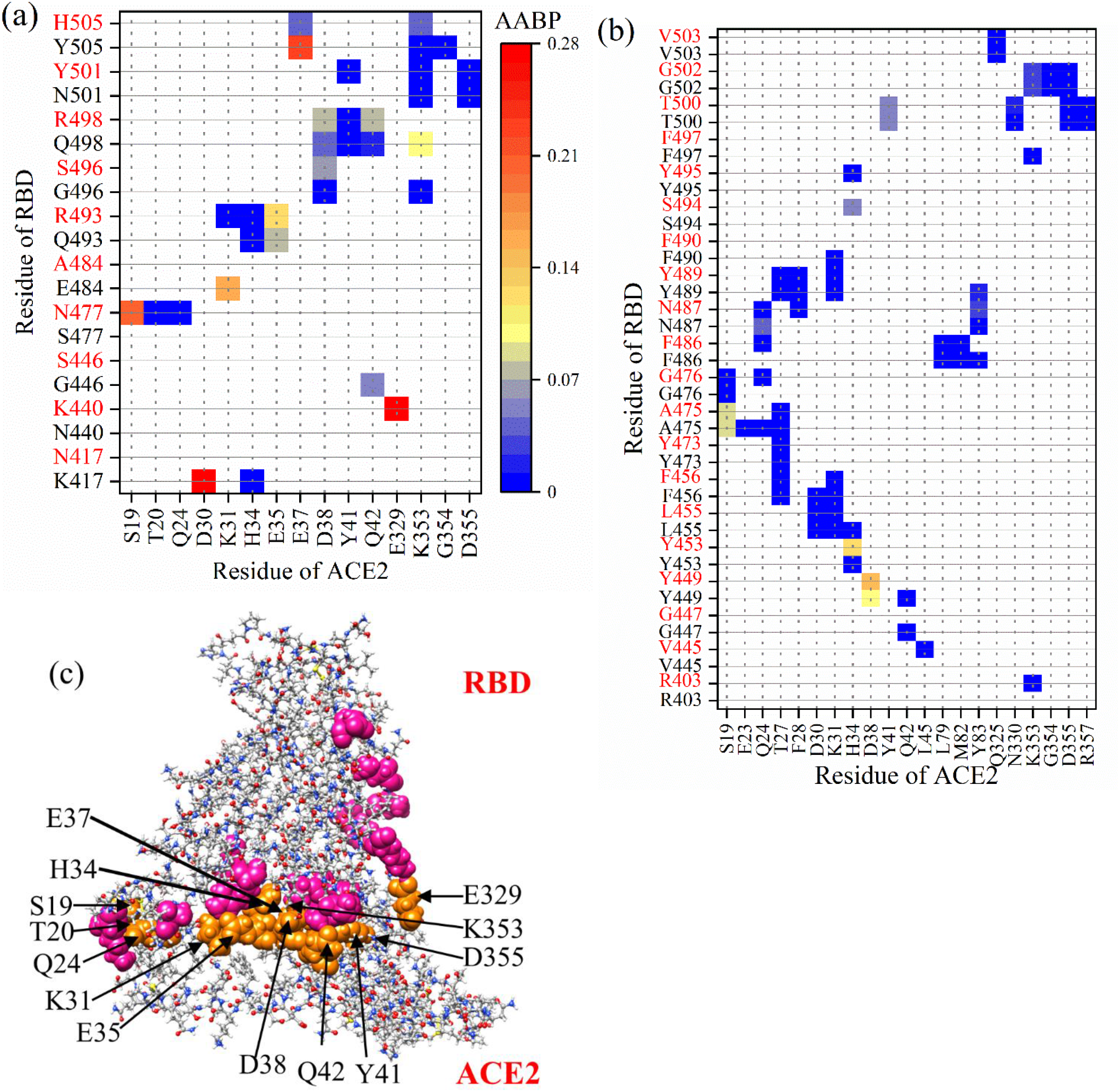
AABP map showing interaction of (a) mutated and (b) unmutated AAs of RBD with ACE2 in the interface model. WT and OV AAs are listed in y-axis labels in black and red color respectively. (c) Ball stick figure of mutated RBD-ACE2 model showing interaction between ACE2 AAs (orange sphere) with mutated AAs of RBD (pink sphere). Grey: C, red: O, blue: N, and white: H

We now analyze the interactions with specific mutations sites. In the case of OV, 7 (K440, N477, R493, S496, R498, Y501, and H505) out of 15 mutated AA interact with ACE2 whereas in the case of WT, 8 (K417, G446, E484, Q493, G496, Q498, N501, and Y505) out of 15 AAs interact with ACE2. Thus, OV interface loses interaction with ACE2 with N417, S446, A484 AAs and gains interaction with K440 and N477. In particular, mutated AAs K440, N477, R493, and S496 shows increase in interaction with ACE2. In our past calculation of interface RBM-ACE2 mutated Y501 showed higher AABP with ACE2 [68], which is consistent with previous findings describing the increase in binding affinity [87-89]. Nevertheless, we note a slight decrease in AABP with ACE2 of Y501 in comparison to N501. In addition, the Y501 interaction with Y41 and D355 has decreased a little in comparison to RBM-ACE2 model [68]. The 7 mutated AAs of RBD interacts with 13 AAs (S19, T20, Q24, K31, H34, E35, E37, D38, Y41, Q42, E329, K353, and D355) of ACE2. Among them, E329 and S19 have strongest bonding with RBD. These 13 AAs are marked in **Figure 10 (c)** clearly showing their proximity to the interface. These AAs can be considered as potential targets since the structure of ACE2 and its interaction with RBD is critical for antibody and drug design [90, 91].

Mutated AAs are definitely the main reason for enhanced interaction between RBD and ACE2. These mutated AAs have an effect not only on the strength of binding with ACE2, but also on the intramolecular interactions of unmutated AAs in the RBD, changing their NN and NL bonding (see **Table 2**). These NN and NL AAs can be both mutated and unmutated ones. Hence, indirectly unmutated AAs also play an important role in changing the overall bonding at the interface. In the RBD, there are 17 unmutated AAs in case of OV and 16 AAs in case of WT that interact with ACE2. The change in interaction of these AAs between WT and OV is also an important point to be considered. OV RBD gains interaction with R403, V445, S494 and loses interaction with F490 and F497.

The AAs in ACE2 interacting with unmutated AAs of RBD are listed in x-axis of **Figure 10 (b)**. Among them E23, T27, F28, D30, L45, L79, M82, Y83, Q325, N330, G354 and R357 only interact with unmutated AAs of RBD. Adding up all these interactions gives the total interface AABP values and shows stronger interaction in OV compared to WT. Hence, these AAs can also be considered as potential target for disruption.

### 4.3 Partial charge and Mechanism of Penetration

As mentioned in **Section 3.4** we have calculated two types of partial charge (PC)—one is for each AAs (PC^AA^), and another is for the entire AABPU unit (PC*). PC* is obtained adding up PC^AA^ of all AAs involved in the AABPU. PC* of AABPU are listed in **Table 2** for 15 sites in RBD for both WT and OV of the interface model. They are also plotted in **Figure 11 (a)**. 10 out of 15 Omicron mutations have changed PC* toward positive values, indicating a significant change in its surface charge distribution that affects either ACE2 binding or antibody binding or both [8, 85].

**Figure 11.**
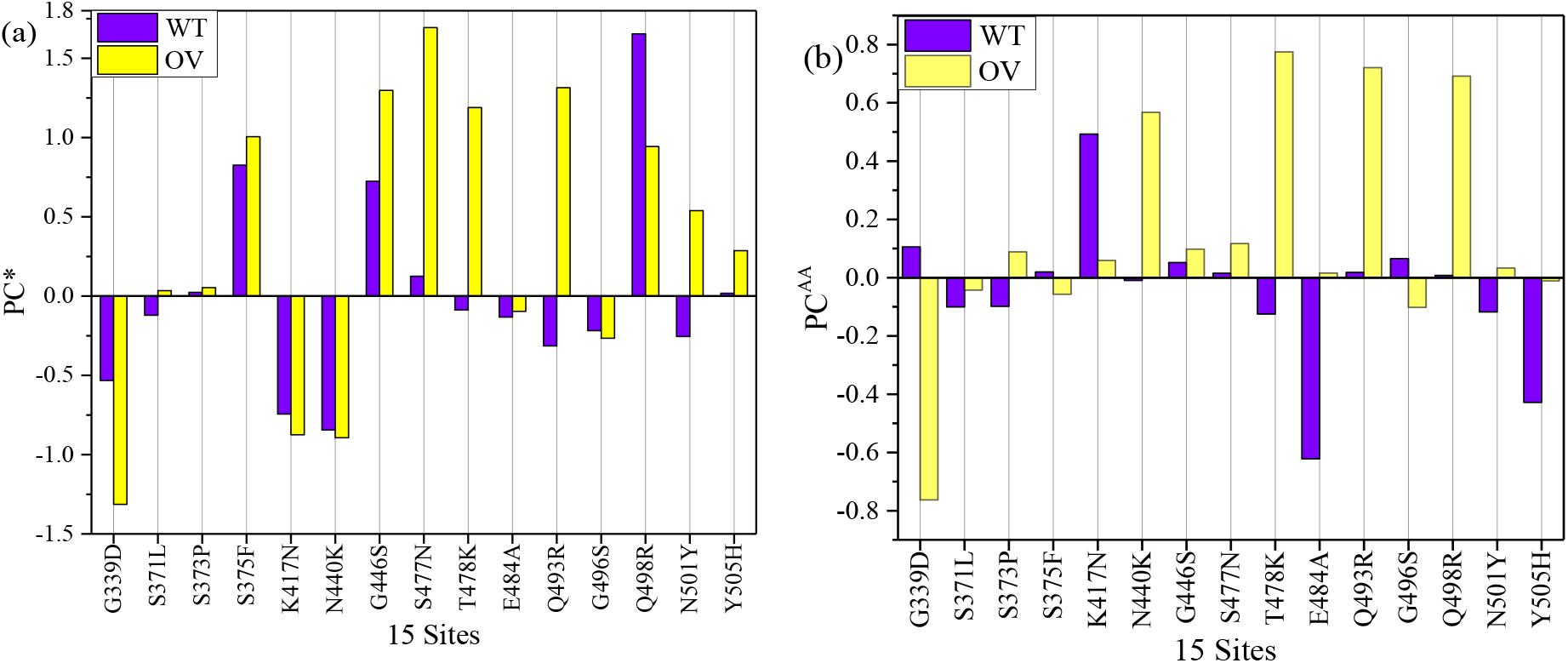
**(a)** Partial Charge per AABPU (PC*) and **(b)** Partial charge per AAs (PC^AA^) for 15 mutation sites in RBD domain of the RBD-ACE2 interface model

**Table 3** shows PC^AA^ for 15 AAs in both WT and OV interface model. They are plotted in **Figure 11 (b)** for easy visualization. Here, the shift toward positive PC is in 11 out of 15 AAs. One of the studies had cited N440K, T478K, Q493R, Q498R, and Y505H to have positive charge [84, 92]. All these AAs falls under the 11 AAs shown in **Figure 11 (b)**. Both PC^AA^ and PC* shows increase in PC in most cases of OV, which is consistent with other studies [93-95]. The sum of PC^AA^ for 15 AAs in WT and OV shown in **Table 3** is -0.727e and 2.185e respectively. Similarly, sum of PC* for 15 AABPU shown in **Table 2** is 0.122e and 4.910e respectively. This is noticeable increase in PC after mutation, which will further increase the electrostatic interaction between RBD and ACE2 or/and RBD and antibodies. This could be one of the major reasons for the rapid infectivity of OV and evade the immunity response from vaccine or other antibody therapeutics.

**Table 3.** Comparison of PC^AA^ between WT and OV of RBD-ACE2

| WT | $PC^{AA}$ | OV | $PC^{AA}$ |
| --- | --- | --- | --- |
| WT G339 | 0.1052 | OV D339 | -0.7636 |
| WT S371 | -0.1003 | OV L371 | -0.0428 |
| WT S373 | -0.0986 | OV P373 | 0.088 |
| WT S375 | 0.0193 | OV F375 | -0.0571 |
| WT K417 | 0.4925 | OV N417 | 0.0586 |
| WT N440 | -0.0097 | OV K440 | 0.5675 |
| WT G446 | 0.0518 | OV S446 | 0.0974 |
| WT S477 | 0.0152 | OV N477 | 0.1168 |
| WT T478 | -0.125 | OV K478 | 0.7745 |
| WT E484 | -0.622 | OV A484 | 0.0153 |
| WT Q493 | 0.0179 | OV R493 | 0.7205 |
| WT G496 | 0.0651 | OV S496 | -0.1027 |
| WT Q498 | 0.0076 | OV R498 | 0.6917 |
| WT N501 | -0.1179 | OV Y501 | 0.0325 |
| WT Y505 | -0.4285 | OV H505 | -0.0115 |

The interacting AAs at the RBD-ACE2 interface can be both mutated and unmutated. We further analyze PC^AA^ values for the AAs having interaction at the interface between RBD (**Figure 12 (a)**) and ACE2 (**Figure 12 (b)**) in WT and OV models. The vertical green lines show common interface interacting AAs in both WT and OV whereas the vertical red lines show the mutated AAs in RBD. There are 5 vertical red lines in **Figure 12 (a)** representing 5 mutated AAs (Q493R, G496S, Q498R, N501Y, and Y505H), which interact with ACE2. Among these 5 mutated AAs 4 have changed PC^AA^ to positive direction, which is consistent with other studies [93-95]. Other unmutated interacting AAs in RBD at the interface have changed PC^AA^ in both positive and negative direction. AAs marked by vertical black lines are interacting in either WT or OV interface. There are 12 such AAs in RBD and 6 of them R403, K440, V445, N477, S494, and Y495 are only interface interacting in OV and remaining 6 AAs K417, G446, G447, E484, F490, and F497 are only interface interacting in WT. In case of ACE2, there are just 5 AAs marked by black vertical lines. T20, L45, and E329 are the only interface interacting in OV and E23 and H34 are the only interface interacting WT. T20, L45, and E329 in OV can be considered as important AAs of ACE2 since they also interact with mutated AAs of RBD. For interacting AAs in ACE2, only F28 and G354 yield a changed PC^AA^ from positive to negative. All remaining AAs have changes in the same direction i.e., either to positive or to negative PC^AA^. This indicates that most of the changes at the interface are due to mutated AAs in RBD.

**Figure 12.**
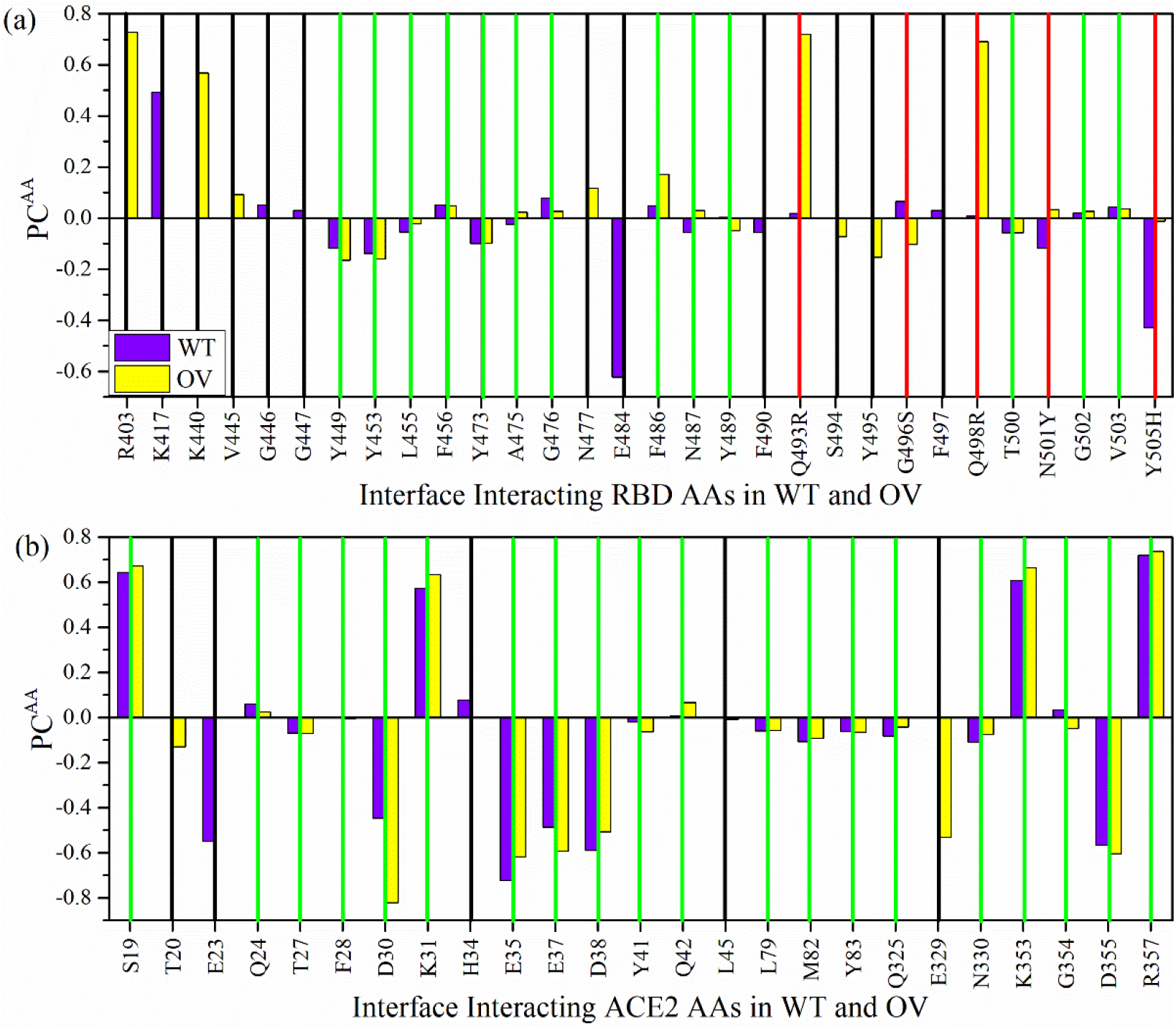
Bar graph for the PC^AA^ of each interacting AA in the interface region of (a) RBD and (b) ACE2 for the WT and OV model. The vertical red lines in (a) denote the mutated AAs in RBD. The vertical green lines denote common interface interacting AAs in both WT and OV. The vertical black lines denote AAs that are interface interacting either in WT or in OV.

In **Figure 13**, we display the PC^AA^ for the RBD-ACE2 interface model in the form of standard solvent excluded layer for both WT ((a), (b), (c), (d)) and OV ((e), (f), (g), (h)). **Figure 13 (c)** and **(g)** shows separated RBD and ACE2, which are further rotated in the in **Figure 13 (d)** and **(h)** with highly positively and negatively charged AA marked. Comparison of PC^AA^ of RBD shows the increase in positive charge in R493, K478, R457, R498 marked in **Figure 13(h)**. Similarly, the PC^AA^ of ACE2 shows the increase in negative charge in E23 and D30, and positive charge in L353 marked in **Figure 13(h)**. PC^AA^ of all AAs in ACE2 for both WT and OV are listed in **Table S2** and **Table S3** respectively. Similarly, PC^AA^ of all AAs in RBD for both WT and OV are listed in **Table S4** and **Table S5** respectively.

**Figure 13.**
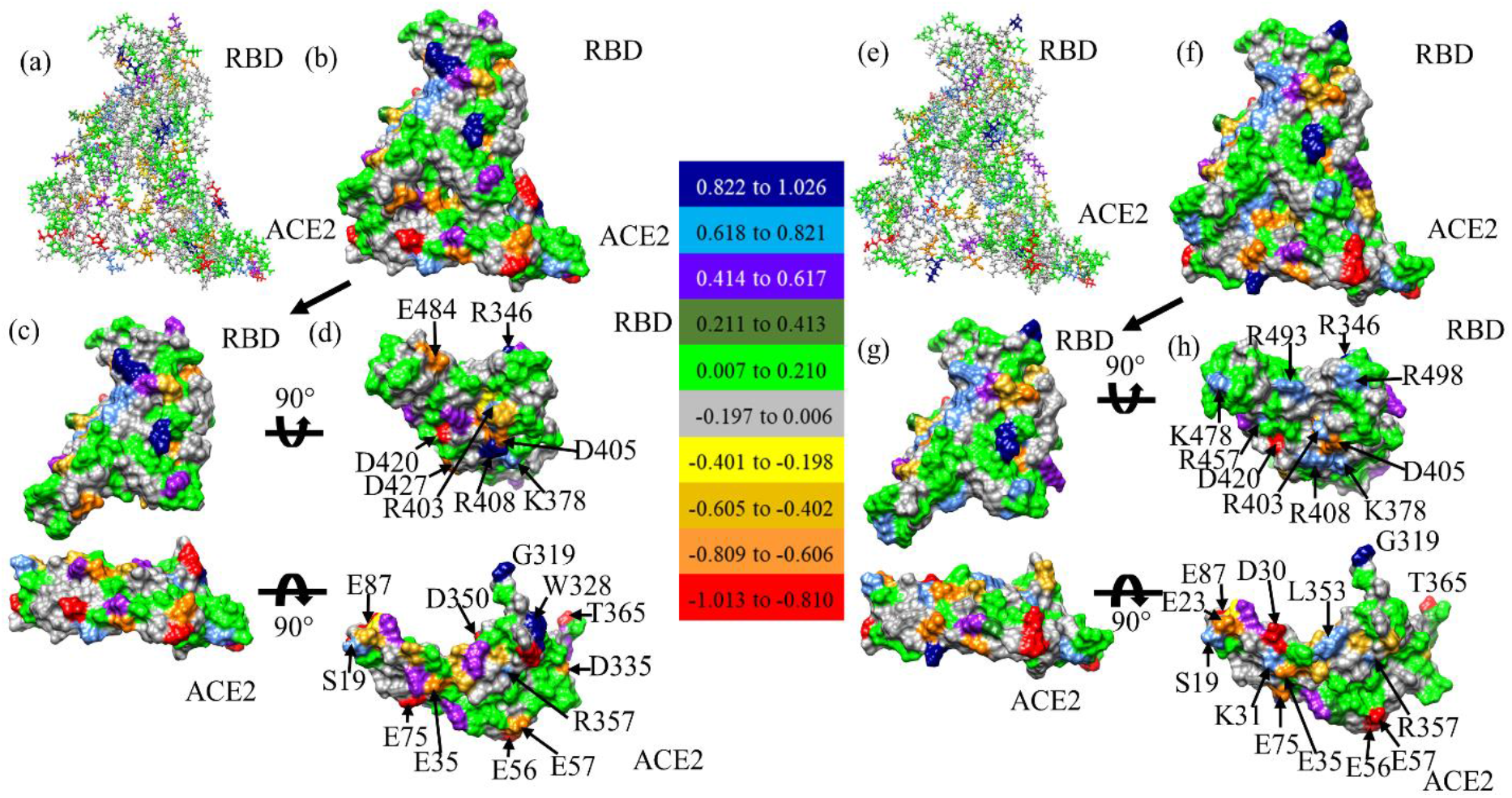
PC^AA^ distribution in the RBD-ACE2 interface in (a) ball and stick, (b) solvent excluded surface, (c) separated surface, and (d) their interacting surface for WT. Similar figures for OV are shown in (e), (f), (g), and (h) respectively. AAs with PC^AA^ higher and lower than 0.618 e and -0.606 e are marked. The color bar shows the total PC^AA^ for different AAs from red (negative) to navy blue (positive).

### 4.4 Implication of RBD-ACE2 Interface on Omicron variants of SARS-CoV-2

It is well known that Omicron variant causes higher infectivity [96]. Based on large-scale *ab initio* calculations we provided some fundamental analysis for 15 mutations in OV and its interaction with ACE2. This includes AABP values indicating the strength of bonding. From AABP, we have analyzed all possible interactions between RBD and ACE2 and have identified all prominent AAs in both RBD and ACE2 that participating in the interaction. Here AABP value predicts the strengthening of bonding between RBD and ACE2. Adding up all AABP values for interacting AAs between RBD-ACE2, we obtained a higher value for OV (1.46e^-^) in comparison to WT (1.33e^-^). Among the mutated AAs K440, N477, R493, and S496 show an increase in binding with ACE2. AAs from ACE2 that interact with mutated and unmutated AAs of RBD have been identified to be S19 and E329 having the strongest bonding with OV RBD.

From the PC calculations, the increase in positive charge in both PC* and PC^AA^ in OV is observed. This dominance in positive charge as a result of mutation is consistent with studies which suggest development of negatively charged antibodies for better binding [93, 95]. Based on PC of interacting AAs, it can be claimed that 80% of the interacting mutated AAs at the interface have changed the partial charge in positive direction. T20, L45, and E329 of ACE2 are the prominent AAs since they interact with RBD in OV whereas this interaction is absent in WT. Prominent changes of PC in the surface of both RBD and ACE2 are identified to be R493, K478, R457, and R498 in RBD and E23 and D30 in ACE2. One of the interesting observations is R493 and R498 with noticeable change of PC in the surface have higher AABP values among 15 mutated AAs. This shows the connection between AABP and PC. The overall increase in volume, increase in sum of AABP between the RBD and ACE2 in OV, and change of PC of most of mutated AAs toward positive charges are important observations which can make OV more lethal and dangerous. Especially, the change in PC after mutation has functional implications as the electrostatic charge modifies its ability to bind strongly with ACE2 or escape for antibody.

## 5.0 Conclusions

In summary, we have provided a detailed account of the mutational effect of Delta and Omicron variants in the SARS-CoV-2 virus, based on large-scale *ab initio* quantum calculations invoking the novel concept of AABPU as a special biomolecular unit. Part 1 is focused on the RBD-SD1 domain and showing the Omicron mutations are much more significant than Delta mutations. We presented the change in the structure of residues involved in mutation including the changes in pertinent hydrogen bonding. Part 2 presents the calculation of the RBD-ACE2 interface complex, showing the much more enhanced binding between RBD and ACE2, providing additional evidence for the increased infectivity of Omicron variants. Specific mutations and their locations at the interface in both RBD and ACE2 are pointed out. We also obtained a detailed partial charge distribution on all the involved AABPUs and their respective central amino acid. These are very valuable data for experimental and clinical scientists. The results of our computations shed additional light on the properties of the emerging VOCs all the way down to the atomic scale.

## Supporting information

Supplementary Materials for the Main

## Supplementary Material

The following supporting information can be down loaded a: https://xxxxxxxxxxxxxxxxxx. Additional descriptions, figures and Tables are provided in the Supplementary Materials.

## Authors’ Contributions

WC, PA and BJ conceived the project. WC and PA performed the calculations. PA and BJ made most of the figures. BJ searched most of the references, WC, BJ, PA drafted the paper together with inputs from RP. All authors participated in the discussion and interpretation of the results. All authors edited and proofread the final manuscript.

## Data Availability Statement

All data are listed in tables or presented in figures in main text or Supplementary Material.

## Acknowledgements

This research used the resources of the National Energy Research Scientific Computing Center supported by DOE under Contract No. DE-AC03-76SF00098 and the Research Computing Support Services (RCSS) of the University of Missouri System. We thank Dr. Richard Gerber, Senior Science Advisor and HPC Department Head for special allocations. This project is funded partly by the National Science Foundation of USA: RAPID DMR/CMMT-2028803 in 2020-2021. RP acknowledges funding from the Key project #12034019 of the National Natural Science Foundation of China.

