## Supplementary Materials for the Main for "Effect of Delta and Omicron mutations on the RBD-SD1 domain of the Spike protein in SARS-CoV-2 and the Omicron mutations on RBD-ACE2 interface complex"

**Content:**

#### S1 Computational Methods

#### (a) VASP

One of the key feature in our computational strategy is to combine the two well established packages based on density functional theory (DFT): The Vienna *ab initio* Simulations package (VASP) [1] and the orthogonalized linear combination of atomic orbital OLCAO) method [2]. This strategy is pivotal for successful application to large and complex materials since more than 20 years ago. In recent years, it has been further demonstrated to be highly effective by using supercells containing large number of atoms. They include but not limited to complex crystals and disordered non-crystalline materials [3-9], biomolecules [10-25], proteins [14, 17-25], glasses [26-30] and different organic [31, 32], inorganic[11, 13, 33] and metallic systems [5, 34-36].

The initial structure for RBD-SD1 domain in S-protein is obtained from the selected source in protein data bank (PDB) with appropriate modification such as addition of missing hydrogen (H) atoms. The initial structure with several thousands of atoms is then placed in a large supercell with periodic boundary conditions. The supercell is sufficiently large to ensure no artificial interaction occurs between the large biomolecule and its periodic image. The model is then fully optimized to high precision using VASP [1], which is known for its efficiency in structural optimization. We adopt the usual projector augmented wave (PAW) method with Perdew-Burke-Ernzerhof (PBE) exchange correlation functional [37] within the generalized gradient approximation (GGA). This selection is one of the several options that balance the accuracy needed and the computational resources available. Detailed tests suggest that the use of the following input parameters to VASP for large biomolecular systems is more than sufficient: (1) energy cut-off at 500 eV; (2) electronic convergence of $10^{-4}$ eV for each step; (3) force convergence for ionic steps at $-10^{-2}$ eV/Å; (4) a single k-point sampling at the center of the supercell (Γ). The final relaxed structure of the supercell models has achieved the accuracy of difference in total energy is less than -0.330 eV or -0.000108 eV per atom. This optimized structure is used as the input data for OLCAO calculation.

#### (b) OLCAO

The OLCAO package also based on DFT was developed by our group at the University of Missouri-Kansas City [2]. It is particularly effective for the calculation of electronic structure and interatomic interactions of large biomolecule systems few other DFT-based methods can match. In contrast to VASP, atomic orbitals are used for basis expansion in conjunction with in orthogonalization to the core orbitals protocol that enable us to diagonalize the huge matrix with a single step to obtain all the energy eigen values and wave functions of the Kohn-Sham equation [38]. Two fundamental quantities from the *ab initio* wave functions from OLCAO are most important: the effective charge ($Q^*$) on each atom and the bond order (BO) value $\rho_{\alpha\beta}$ between any pair atoms α and β in the supercell defined in Eq. (1) and (2) below.

$$Q_\alpha^* = \sum_i \sum_{m,occ} \sum_{j,\beta} C_{i\alpha}^{*m} C_{j\beta}^m S_{i\alpha,j\beta} \quad (1)$$

$$\rho_{\alpha\beta} = \sum_{m,occ} \sum_{i,j} C_{i\alpha}^{*m} C_{j\beta}^m S_{i\alpha,j\beta} \quad (2)$$

In equations (1) and (2), $S_{i\alpha,j\beta}$ are the overlap integrals between the $i^{th}$ orbital in the $\alpha^{th}$ atom and the $j^{th}$ orbital in the $\beta^{th}$ atom. $C_{j\beta}^m$ are the eigenvector coefficients of the $m^{th}$occupied molecular orbital. We can the so-called partial charge (PC) from $Q_\alpha^*$ , which is the deviation of $Q_\alpha^*$ from the neutral atomic charge $Q_\alpha^0$ on the same atom ($\Delta Q_\alpha = Q_\alpha^0 - Q_\alpha^*$ ). The BO represents the strength of the bond between two atoms in the unit of electrons ($e^-$). BO usually scales with the bond length (BL) or the distance of separation between atoms α and β, depending also on the local atomic configuration of the vicinal atoms. It should be explicitly pointed out that PC and BO in Eq. (1) and (2) are fundamentally different form other simulation methods which are fixed parameters such as in the force field specification in molecular dynamic (MD) simulation. Another point to emphasize is that Q* and BO are basis-dependent since they are based on the Mulliken scheme [39, 40] using localized atomic orbitals. We use the minimal basis in most of our calculations for large biomolecular systems. Another key important point is that OLCAO is a one-point calculation to obtain all BO values for all atomic pairs to characterize the internal cohesion of the system under study, rather than the traditional total energy or enthalpy calculation (two-point or even many-point calculations) which is used to describe the strength of binding between biomolecular systems. The sum of all BO values within a structural component such as in a protein and its subdomains gives the total bond order (TBO), which accurately describes the internal cohesion critical to analyzing the AA-AA network for large complex biomolecules.

**(c) Analysis of Amino acid - amino acid bond pair unit (AABPU)**

In complex biomolecules we need to extend the concept of the bond order (BO) values for a *pair of atoms* to interaction between a *pair of amino acids* (AAs). We refer to this generalized quantifier of molecular interactions the *amino acid-amino acid bond pair* (AABP) as first described in ref. [17]

$$AABP(u,v) = \sum_{\alpha \epsilon u} \sum_{\beta \epsilon v} \rho_{\alpha i,\beta j} \quad (3)$$

In Eq. (3), the summations are over all atoms $\alpha$ in $AA\ u$ and all atoms $\beta$ in $AA\ v$. AABP considers all possible bonding between two AAs including both covalent and hydrogen bonding (HB). AABP value is a single parameter proxy that quantifies the interaction between two AAs. The stronger the interaction, the higher will be the AABP value and vice versa irrespective of the nature and composition of the 20 canonical AAs. The specific structural unit that contains the relevant AAs is coined as AABPU. AABP value in each AABPU can be further resolved into different components, nearest neighbor (NN) in the amino acid sequence (NN-AAPB) and non-local (NL-AAPB) parts. It should be emphasized that the AABP does not involve the "BL" used for the description of interacting atoms since the distance of separation between two AAs is impossible to quantify precisely even people have been tried by using distance of separation between specific "C" atom in the AAs. AABP values are calculated from quantum mechanical wave functions of the entire biomolecular unit and thus represent a collective structural parameter, including the effects of all atomic pairs involved. AABPU is a novel concept to measure of molecular interactions in biomolecules, that contains the nearest-neighbor or local interactions of AAs that are vicinal along the sequence and in the 3D folding space, as well as the off-diagonal or non-local interaction between AAs that are not vicinal in the sequence space but are interacting in 3D folding space. Clearly, this is a giant step forward in in the theory of biomolecular interaction.

**(d) Graphical Illustration**

The methods used for graphical illustrations are briefly outlined below: **Figure 1 (a)** is prepared using the PowerPoint whereas **Figure 1 (b)**, **(c)**, **(d)** and **Figures 2**, **3,** and **13** are prepared using Chimera [41]. All other figures (**Figures 4** to **Figure 12)** are prepared using Origin-version 8 software. The graphical illustration of AABPU in **Figure 3** and **Figure S1** entails the plots of a 3-dimensional (3D) structure of a collection of AAs on a 2D plane in atomic scale. This is a complex and time-consuming task. We proceed it as follows: Firstly, the data are from optimized structure using VASP followed by OLCAO calculations. The numerical data for the bonding between every pair of bonds are extracted are listed in a large table. These bonds are then analyzed in different groups such as bond types, (covalent or hydrogen bond), bonds formed by specific amino acids, mutated or unmutated (WT) etc. All amino acids involved in bonding with the mutated amino acids are considered. We then prepare the plots for the complicated 2D plane figure via Chimera [41].

**S2 Partial density of states (PDOS) for WT, DV and OV in RBD-SD1**

In **Figure S3**, we display the 18 PDOS following the order in **Table 1** with WT and DV or OV in the same figure. To simplify the discussion, the structural unit used in PDOS are the central amino acid for each model listed in **Table 1**. The following interesting observations are noted

A. PDOS for all panels in **Figure S3** are very close between WT, but the mutated types show many differences. This demonstrates the penetrating details can be revealed in PDOS.
B. The peak positions in PDOS are mostly similar and aligned, a fact consistent with all biological molecules consists of AAs.
C. The PDOS figures in **Figure S3** are for each main AA in **Table 1**. They should not be compared or correlated with AABPU since the later consists of additional NN, Nonlocal amino acids with contributions from HBs.
Still, for a single AA in the AABPU, some interesting observations can be identified. We will comment on each of the 18 AAs.
D. The comparative study is based on following observations: increase or decrease in the area under the curve which is the number of energy states it contains.
E. There are roughly two regions in the VB, the first major peak between 0 to -4 eV and the second dominating group of states below -4 eV.
F. In the unoccupied CB regions, they can also be roughly divided into a lower peak below 5 eV and all other states above it. They are the antibonding images of the states in the VB groups.

We succinctly comment on each PDOS figure by comparing the observed features in different regions before and after mutation.
1. DV L452: In both the VB and CB, the areas under the PDOS curves are increased after mutation.
2. DV K478: In the VB, both areas under the curves are increased after mutation except the first peak remain similar. The features in the CB are similar as the mirror image of the VB.
3. OV D339: In both the VB and CB, the areas under the PDOS curves are significantly increased after mutation.
4. OV L371: In both the VB and CB, the areas under the PDOS curves are increased after mutation except the trend is reversed in the first peak compared to the second one. The increase is larger in the second peak in CB than in the VB.
5. OV P373: The feature in both VB and CB are similar to OV L371 except the change after mutation is by a lesser amount.

6. OV F375: The feature in both VB and CB are similar to OV P373 except the change after mutation is by a larger amount.
7. OV N417: This figure is the first one in which the area under the curve in both VB and CB are decreased by a fairly large amount after mutation, especially in the second peak in the CB.
8. OV K440: The feature in both VB and CB are similar to OV P373 except the change after mutation is by a larger amount in the second peak of the CB.
9. OV S446: Both areas under the curve in VB and CB increased after mutation similar to K440.
10. OV N477: This site is very similar to OV N417 in almost all aspects. This is the second case where mutation resulted in the areas under the curve.
11. OV K478: This site is very similar to OV K440 in almost all aspects.
12. OV A484: This site is very similar to OV N477 in almost all aspects. This is the third case where mutation reduces the areas under the curve
13. OV R493: This site is very similar to OV K478 in almost all aspects.
14. OV S496: This site is almost identical to OV N477 in shapes of the curve but with very puzzling difference. Mutation in increases the area under the curves whereas OV N477 is opposite.
15. OV R498: This site is very similar to OV R493 in almost all aspects.
16. OV Y501: This site is again similar to OV R493 as well except the increase in areas under curves after mutation is slightly larger.
17. OV H505: This site is very similar to OV A484 in almost all aspects. This is the last example where mutation actually decreases the areas under the curves.
18. OV K547: This site is very similar to OV K493 in almost all aspects. We emphasize that this is the only site in the SD1 portion of the structural model, not the RBD.
19. The two DV sites have very similar PDOS spectra. Mutation increases the areas under the curves.
20. Out of 16 OV sites, 4 of them have mutation reduces the areas under the curve (25%). OV sites also have more variations among them such as difference between VB and CB regions. These observations all point to the complexity of Omicron variant in addition to large number of mutations.

### Supplementary Figures:

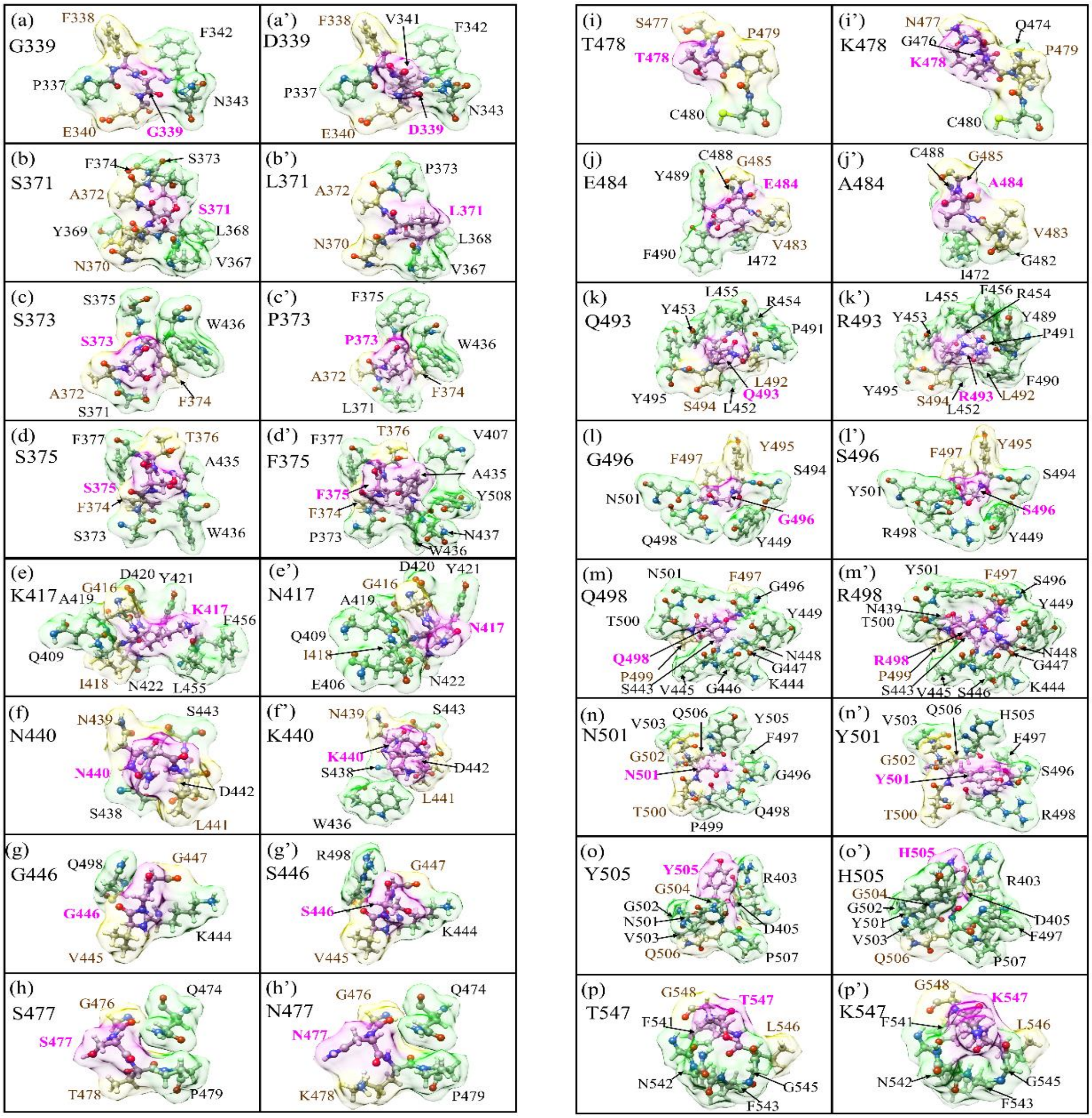


**Figure S1.** Details of the shape change of AABPU of the sixteen mutation sites in RBD-SD1: (a) G339, (b) S371, (c) S373, (d) S375, (e) K417, (f) N440, (g) G446, (h) S477, (i) T478, (j) E484, (k) Q493, (l) G496, (m) Q498, (n) N501, (o) Y505, and (p) T547 for the WT. (a') D339, (b') L371, (c') P373, (d') F375, (e') N417, (f') K440, (g') S446, (h') N477, (i') K478, (j') A484, (k') R493, (l') S496, (m') R498, (n') Y501, (o') H505, and (p') K547 for the OV. The surface of mutated sites is shown in magenta, surface of NN and NL are shown in yellow and green respectively. All NN and NL AAs are marked near to their surface in brown and black respectively.

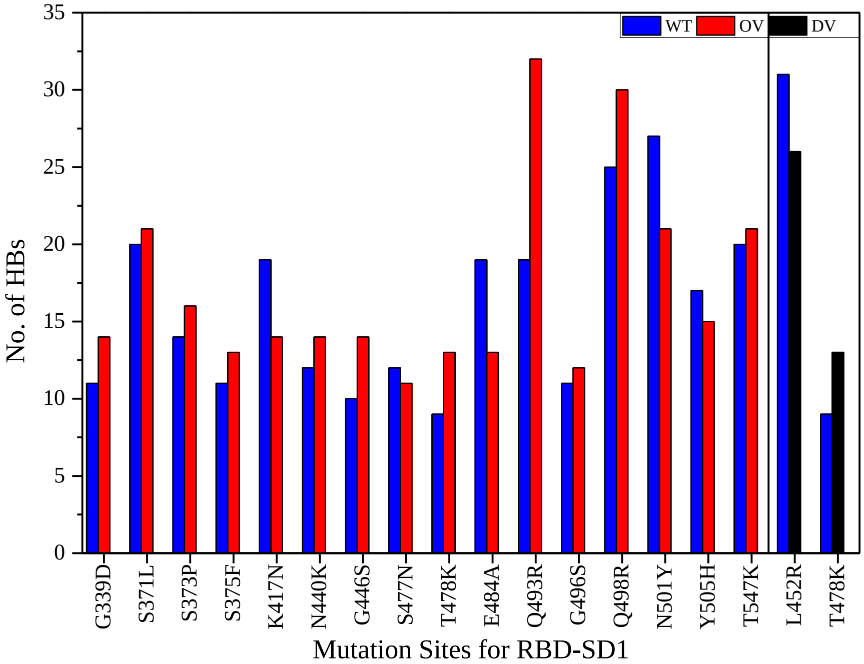


**Figure S2.** Number of hydrogen bonds in 18 mutation sites of RBD-SD1 including WT, DV, and OV.

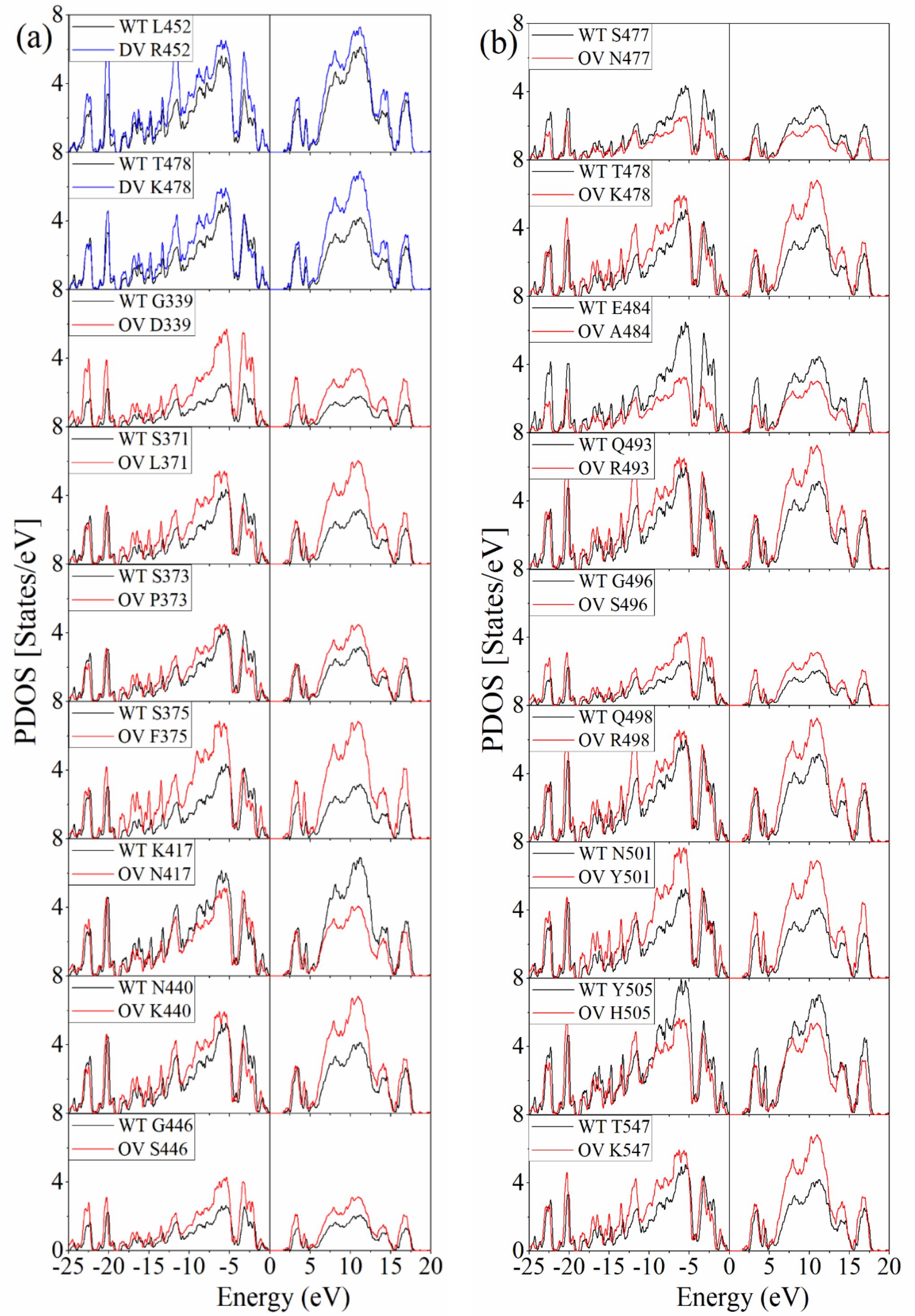


**Figure S3.** Comparison of PDOS per each amino acid for the 18 mutations (16 for OV, 2 for DV) with the WT. Black: WT, blue: DV, red: OV. For easy contrast, each panel for the listed AA site has two PDOS curves, WT and mutated one.

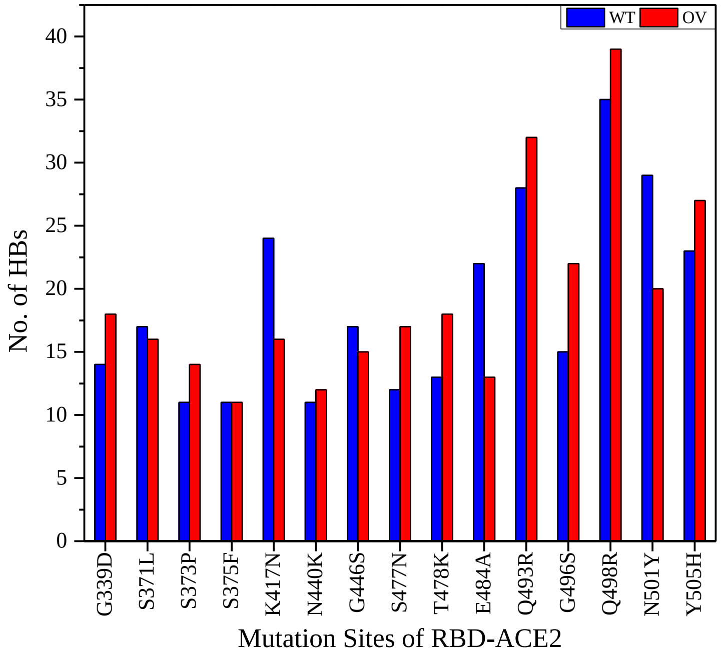


**Figure S4.** Number of hydrogen bonds in 15 mutation sites of RBD-ACE2 including WT and OV.

### Supplementary Tables:

**Table S1:** Number of HBs and total bond order (TBO) in RBD-SD1 for WT, DV, and OV shown in **Figure 6**. The last column is the total data points for HB.

| **HBs** | | **O…H** | **N…H** | **Total** |
|---|---|---|---|---|
| **WT** | Count (BL<2.0Å) | 133 | 4 | |
| | Count (BL>2.0Å) | 2391 | 1493 | |
| | Count (Total) | 2524 | 1497 | 4021 |
| | TBO <2.0Å | 6.2467 | 0.1819 | |
| | TBO >2.0Å | 7.7355 | 3.0292 | |
| | TBO (Total) | 13.9822 | 3.2111 | 17.1933 |
| **DV** | Count (BL<2.0Å) | 133 | 4 | |
| | Count (BL>2.0Å) | 2384 | 1516 | |
| | Count (Total) | 2517 | 1520 | 4037 |
| | TBO <2.0Å | 6.3117 | 0.1807 | |
| | TBO >2.0Å | 7.6313 | 3.1029 | |
| | TBO (Total) | 13.943 | 3.2836 | 17.2266 |
| **OV** | Count (BL<2.0Å) | 146 | 5 | |
| | Count (BL>2.0Å) | 2384 | 1549 | |
| | Count (Total) | 2530 | 1554 | 4084 |
| | TBO <2.0Å | 6.7742 | 0.3539 | |
| | TBO >2.0Å | 7.4242 | 3.2033 | |
| | TBO (Total) | 14.1984 | 3.5572 | 17.7556 |

**Table S2**: $PC^{AA}$ for ACE2 AAs of WT model. Color coded according to **Figure 13**. The residues in ACE2: 19-88 and 319-365.

| AAs | $PC^{AA}$ | AAs | $PC^{AA}$ | AAs | $PC^{AA}$ | AAs | $PC^{AA}$ |
|---|---|---|---|---|---|---|---|
| S19 | 0.644 | W48 | 0.025 | S77 | -0.015 | P336 | 0.047 |
| T20 | -0.118 | N49 | 0.045 | T78 | -0.030 | G337 | -0.027 |
| I21 | 0.023 | Y50 | 0.030 | L79 | -0.060 | N338 | 0.120 |
| E22 | -0.393 | N51 | -0.071 | A80 | 0.002 | V339 | 0.057 |
| E23 | -0.549 | T52 | -0.067 | Q81 | -0.016 | Q340 | 0.017 |
| Q24 | 0.061 | N53 | 0.036 | M82 | -0.108 | K341 | 0.657 |
| A25 | 0.002 | I54 | 0.017 | Y83 | -0.061 | A342 | 0.042 |
| K26 | 0.512 | T55 | -0.157 | P84 | 0.057 | V343 | -0.089 |
| T27 | -0.070 | E56 | -0.815 | L85 | 0.135 | C344 | 0.123 |
| F28 | 0.002 | E57 | -0.780 | Q86 | -0.105 | H345 | -0.132 |
| L29 | 0.040 | N58 | -0.034 | E87 | -1.010 | P346 | 0.144 |
| D30 | -0.446 | V59 | 0.010 | I88 | -0.998 | T347 | -0.101 |
| K31 | 0.573 | Q60 | -0.045 | G319 | 1.031 | A348 | 0.048 |
| F32 | -0.081 | N61 | 0.053 | L320 | -0.184 | W349 | 1.898 |
| N33 | -0.016 | M62 | -0.034 | P321 | 0.162 | D350 | -0.814 |
| H34 | 0.078 | N63 | -0.016 | N322 | -0.034 | L351 | -0.089 |
| E35 | -0.723 | N64 | 0.198 | M323 | 0.005 | G352 | -0.032 |
| A36 | -0.084 | A65 | 0.097 | T324 | 0.045 | K353 | 0.607 |
| E37 | -0.486 | G66 | -0.038 | Q325 | -0.081 | G354 | 0.033 |
| D38 | -0.589 | D67 | -0.774 | G326 | 0.040 | D355 | -0.567 |
| L39 | 0.027 | K68 | 0.519 | F327 | 0.051 | F356 | -0.034 |
| F40 | -0.008 | W69 | 0.049 | W328 | 2.092 | R357 | 0.719 |
| Y41 | -0.020 | S70 | -0.021 | E329 | -0.882 | I358 | 0.027 |
| Q42 | 0.006 | A71 | 0.076 | N330 | -0.109 | L359 | -0.048 |
| S43 | -0.006 | F72 | -0.009 | S331 | -0.079 | M360 | -0.027 |
| S44 | -0.036 | L73 | -0.001 | M332 | 0.045 | C361 | 0.082 |
| L45 | -0.016 | K74 | 0.816 | L333 | 0.021 | T362 | -0.057 |
| A46 | 0.048 | E75 | -0.929 | T334 | 0.093 | K363 | 0.498 |
| S47 | -0.049 | Q76 | -0.006 | D335 | -0.621 | V364 | 0.026 |
| | | | | | | T365 | -1.006 |

**Table S3**: $PC^{AA}$ for ACE2 AAs of OV model. Color coded according to **Figure 13**. The residues in ACE2: 19-88 and 319-365.

| **AAs** | **$PC^{AA}$** | **AAs** | **$PC^{AA}$** | **AAs** | **$PC^{AA}$** | **AAs** | **$PC^{AA}$** |
|---|---|---|---|---|---|---|---|
| S19 | 0.672 | W48 | 0.028 | S77 | -0.016 | P336 | 0.050 |
| T20 | -0.131 | N49 | 0.027 | T78 | -0.066 | G337 | 0.006 |
| I21 | -0.004 | Y50 | 0.033 | L79 | -0.057 | N338 | 0.126 |
| E22 | -0.399 | N51 | -0.057 | A80 | -0.019 | V339 | 0.049 |
| E23 | -0.670 | T52 | -0.063 | Q81 | 0.028 | Q340 | -0.070 |
| Q24 | 0.024 | N53 | -0.003 | M82 | -0.092 | K341 | 0.718 |
| A25 | 0.002 | I54 | 0.004 | Y83 | -0.065 | A342 | 0.041 |
| K26 | 0.505 | T55 | -0.066 | P84 | 0.077 | V343 | -0.084 |
| T27 | -0.071 | E56 | -0.828 | L85 | 0.161 | C344 | 0.136 |
| F28 | -0.005 | E57 | -0.918 | Q86 | -0.165 | H345 | -0.127 |
| L29 | -0.020 | N58 | 0.080 | E87 | -0.953 | P346 | 0.094 |
| D30 | -0.821 | V59 | 0.006 | I88 | -1.013 | T347 | -0.033 |
| K31 | 0.634 | Q60 | -0.130 | G319 | 1.026 | A348 | 0.028 |
| F32 | -0.074 | N61 | 0.046 | L320 | -0.146 | W349 | 1.912 |
| N33 | -0.072 | M62 | -0.039 | P321 | 0.135 | D350 | -0.707 |
| H34 | 0.161 | N63 | -0.033 | N322 | -0.033 | L351 | -0.052 |
| E35 | -0.617 | N64 | 0.239 | M323 | -0.001 | G352 | -0.075 |
| A36 | -0.050 | A65 | 0.098 | T324 | 0.059 | K353 | 0.664 |
| E37 | -0.594 | G66 | -0.031 | Q325 | -0.041 | G354 | -0.048 |
| D38 | -0.508 | D67 | -0.796 | G326 | 0.060 | D355 | -0.604 |
| L39 | -0.022 | K68 | 0.606 | F327 | 0.024 | F356 | -0.032 |
| F40 | -0.010 | W69 | -0.004 | W328 | 0.101 | R357 | 0.736 |
| Y41 | -0.064 | S70 | -0.027 | E329 | -0.531 | I358 | 0.033 |
| Q42 | 0.066 | A71 | -0.026 | N330 | -0.075 | L359 | -0.062 |
| S43 | 0.027 | F72 | 0.022 | S331 | -0.072 | M360 | -0.001 |
| S44 | -0.047 | L73 | 0.008 | M332 | 0.033 | C361 | 0.048 |
| L45 | -0.009 | K74 | 0.990 | L333 | 0.002 | T362 | -0.078 |
| A46 | 0.074 | E75 | -0.808 | T334 | 0.057 | K363 | 0.401 |
| S47 | -0.044 | Q76 | -0.059 | D335 | -0.471 | V364 | 0.013 |
| | | | | | | T365 | -0.992 |

| **Table S4**: PC$^{AA}$ for RBD AAs of WT model. Color coded according to **Figure 13**. The residues in RBD: 333-526. | | | | | | | |
|---|---|---|---|---|---|---|---|
| **AAs** | **PC$^{AA}$** | **AAs** | **PC$^{AA}$** | **AAs** | **PC$^{AA}$** | **AAs** | **PC$^{AA}$** |
| T333 | 0.510 | V382 | -0.022 | G431 | -0.040 | C480 | -0.078 |
| N334 | -0.090 | S383 | -0.106 | C432 | 0.079 | N481 | 0.074 |
| L335 | 0.159 | P384 | 0.171 | V433 | -0.036 | G482 | -0.065 |
| C336 | -0.193 | T385 | 0.050 | I434 | -0.014 | V483 | -0.020 |
| P337 | 0.195 | K386 | 0.448 | A435 | -0.033 | E484 | -0.622 |
| F338 | -0.069 | L387 | -0.038 | W436 | 0.017 | G485 | -0.074 |
| G339 | 0.105 | N388 | -0.010 | N437 | 0.079 | F486 | 0.048 |
| E340 | -0.458 | D389 | -0.482 | S438 | -0.225 | N487 | -0.056 |
| V341 | -0.063 | L390 | 0.001 | N439 | 0.072 | C488 | 0.064 |
| F342 | 0.001 | C391 | 0.008 | N440 | -0.010 | Y489 | 0.003 |
| N343 | -0.050 | F392 | -0.021 | L441 | -0.008 | F490 | -0.056 |
| A344 | 0.007 | T393 | -0.064 | D442 | -0.675 | P491 | 0.019 |
| T345 | 0.036 | N394 | 0.077 | S443 | 0.027 | L492 | 0.066 |
| R346 | 0.866 | V395 | 0.017 | K444 | 0.613 | Q493 | 0.018 |
| F347 | 0.006 | Y396 | -0.173 | V445 | 0.132 | S494 | -0.043 |
| A348 | 0.005 | A397 | 0.005 | G446 | 0.052 | Y495 | -0.213 |
| S349 | 0.050 | D398 | -0.647 | G447 | 0.030 | G496 | 0.065 |
| V350 | -0.007 | S399 | 0.063 | N448 | 0.037 | F497 | 0.029 |
| Y351 | 0.064 | F400 | 0.016 | Y449 | -0.117 | Q498 | 0.008 |
| A352 | -0.051 | V401 | 0.024 | N450 | 0.160 | P499 | 0.178 |
| W353 | -0.017 | I402 | -0.048 | Y451 | -0.101 | T500 | -0.058 |
| N354 | -0.006 | R403 | 0.816 | L452 | -0.010 | N501 | -0.118 |
| R355 | 0.720 | G404 | 0.041 | Y453 | -0.139 | G502 | 0.020 |
| K356 | 0.458 | D405 | -0.702 | R454 | 0.687 | V503 | 0.043 |
| R357 | 0.914 | E406 | -0.759 | L455 | -0.054 | G504 | 0.098 |
| I358 | -0.132 | V407 | -0.034 | F456 | 0.052 | Y505 | -0.429 |
| S359 | -0.046 | R408 | 0.828 | R457 | 0.753 | Q506 | -0.033 |
| N360 | -0.028 | Q409 | -0.013 | K458 | 0.551 | P507 | 0.140 |
| C361 | 0.048 | I410 | 0.003 | S459 | -0.113 | Y508 | -0.024 |
| V362 | -0.112 | A411 | -0.106 | N460 | 0.047 | R509 | 0.683 |
| A363 | 0.031 | P412 | 0.144 | L461 | 0.014 | V510 | 0.000 |
| D364 | -0.651 | G413 | -0.040 | K462 | 0.397 | V511 | -0.076 |
| Y365 | -0.093 | Q414 | -0.014 | P463 | 0.128 | V512 | 0.012 |
| S366 | -0.101 | T415 | 0.022 | F464 | -0.052 | L513 | -0.007 |
| V367 | -0.027 | G416 | -0.078 | E465 | -0.433 | S514 | 0.001 |
| L368 | -0.032 | K417 | 0.493 | R466 | 0.784 | F515 | 0.093 |
| Y369 | 0.003 | I418 | -0.045 | D467 | -0.598 | E516 | -0.646 |
| N370 | -0.053 | A419 | 0.103 | I468 | 0.037 | L517 | 0.004 |
| S371 | -0.100 | D420 | -0.832 | S469 | -0.130 | L518 | -0.031 |
| A372 | 0.146 | Y421 | 0.031 | T470 | 0.079 | H519 | 0.078 |
| S373 | -0.099 | N422 | -0.034 | E471 | -0.508 | A520 | -0.109 |
| F374 | 0.041 | Y423 | -0.136 | I472 | -0.005 | P521 | 0.120 |
| S375 | 0.019 | K424 | 0.660 | Y473 | -0.100 | A522 | 0.076 |
| T376 | 0.038 | L425 | -0.013 | Q474 | 0.024 | T523 | -0.112 |
| F377 | 0.047 | P426 | 0.084 | A475 | -0.024 | V524 | -0.044 |
| K378 | 0.819 | D427 | -0.628 | G476 | 0.079 | C525 | 0.069 |
| C379 | 0.027 | D428 | -0.986 | S477 | 0.015 | G526 | -0.483 |
| Y380 | -0.035 | F429 | -0.016 | T478 | -0.125 | | |
| G381 | 0.026 | T430 | -0.037 | P479 | 0.132 | | |

**Table S5**: $PC^{AA}$ for RBD AAs of OV model. Color coded according to **Figure 13**. The residues in RBD: 333-526.

| AAs | $PC^{AA}$ | AAs | $PC^{AA}$ | AAs | $PC^{AA}$ | AAs | $PC^{AA}$ |
|---|---|---|---|---|---|---|---|
| T333 | 0.914 | V382 | -0.067 | G431 | -0.046 | C480 | -0.110 |
| N334 | -0.039 | S383 | -0.019 | C432 | 0.106 | N481 | 0.056 |
| L335 | 0.093 | P384 | 0.040 | V433 | -0.062 | G482 | -0.005 |
| C336 | -0.172 | T385 | -0.006 | I434 | -0.009 | V483 | -0.035 |
| P337 | 0.173 | K386 | 0.513 | A435 | 0.011 | A484 | 0.015 |
| F338 | -0.077 | L387 | 0.084 | W436 | 0.057 | G485 | -0.087 |
| D339 | -0.764 | N388 | -0.150 | N437 | 0.095 | F486 | 0.170 |
| E340 | -0.442 | D389 | -0.923 | S438 | -0.231 | N487 | 0.030 |
| V341 | -0.053 | L390 | -0.017 | N439 | 0.059 | C488 | 0.031 |
| F342 | -0.011 | C391 | 0.000 | K440 | 0.568 | Y489 | -0.049 |
| N343 | -0.130 | F392 | -0.046 | L441 | -0.025 | F490 | 0.026 |
| A344 | -0.010 | T393 | -0.011 | D442 | -0.734 | P491 | 0.159 |
| T345 | 0.011 | N394 | 0.134 | S443 | 0.032 | L492 | 0.007 |
| R346 | 0.949 | V395 | -0.013 | K444 | 0.647 | R493 | 0.721 |
| F347 | 0.032 | Y396 | -0.165 | V445 | 0.092 | S494 | -0.073 |
| A348 | 0.082 | A397 | 0.019 | S446 | 0.097 | Y495 | -0.153 |
| S349 | 0.055 | D398 | -0.646 | G447 | -0.066 | S496 | -0.103 |
| V350 | 0.002 | S399 | 0.051 | N448 | 0.047 | F497 | 0.030 |
| Y351 | 0.072 | F400 | 0.010 | Y449 | -0.165 | R498 | 0.692 |
| A352 | -0.067 | V401 | 0.035 | N450 | 0.153 | P499 | 0.112 |
| W353 | -0.005 | I402 | -0.045 | Y451 | -0.114 | T500 | -0.058 |
| N354 | -0.001 | R403 | 0.728 | L452 | 0.001 | Y501 | 0.032 |
| R355 | 0.713 | G404 | 0.005 | Y453 | -0.160 | G502 | 0.026 |
| K356 | 0.434 | D405 | -0.757 | R454 | 0.680 | V503 | 0.036 |
| R357 | 0.795 | E406 | -0.779 | L455 | -0.022 | G504 | 0.094 |
| I358 | -0.095 | V407 | -0.009 | F456 | 0.049 | H505 | -0.012 |
| S359 | -0.009 | R408 | 0.786 | R457 | 0.791 | Q506 | -0.101 |
| N360 | 0.027 | Q409 | 0.002 | K458 | 0.546 | P507 | 0.141 |
| C361 | 0.138 | I410 | 0.037 | S459 | -0.131 | Y508 | -0.063 |
| V362 | -0.080 | A411 | -0.099 | N460 | 0.005 | R509 | 0.652 |
| A363 | -0.073 | P412 | 0.131 | L461 | 0.009 | V510 | -0.007 |
| D364 | -0.526 | G413 | -0.066 | K462 | 0.402 | V511 | -0.055 |
| Y365 | 0.059 | Q414 | 0.071 | P463 | 0.137 | V512 | -0.005 |
| S366 | 0.064 | T415 | -0.014 | F464 | -0.053 | L513 | 0.001 |
| V367 | -0.013 | G416 | -0.039 | E465 | -0.463 | S514 | -0.022 |
| L368 | 0.036 | N417 | 0.059 | R466 | 0.799 | F515 | 0.094 |
| Y369 | -0.096 | I418 | -0.018 | D467 | -0.676 | E516 | -0.692 |
| N370 | -0.005 | A419 | 0.073 | I468 | 0.060 | L517 | 0.072 |
| L371 | -0.043 | D420 | -0.811 | S469 | 0.004 | L518 | -0.039 |
| A372 | -0.083 | Y421 | 0.036 | T470 | 0.064 | H519 | 0.069 |
| P373 | 0.088 | N422 | -0.043 | E471 | -0.541 | A520 | -0.144 |
| F374 | 0.091 | Y423 | -0.125 | I472 | -0.021 | P521 | 0.138 |
| F375 | -0.057 | K424 | 0.631 | Y473 | -0.098 | A522 | 0.025 |
| T376 | -0.016 | L425 | 0.013 | Q474 | 0.092 | T523 | -0.157 |
| F377 | 0.066 | P426 | 0.138 | A475 | 0.024 | V524 | -0.040 |
| K378 | 0.800 | D427 | -0.558 | G476 | 0.026 | C525 | 0.066 |
| C379 | 0.050 | D428 | -0.949 | N477 | 0.117 | G526 | -0.813 |
| Y380 | -0.179 | F429 | -0.039 | K478 | 0.775 | | |
| G381 | 0.175 | T430 | -0.048 | P479 | 0.120 | | |
